# A hundred genes implicated in intellectual disability and autism regulate habituation learning and reveal an opposing role for Ras-MAPK signaling in inhibitory and excitatory neurons

**DOI:** 10.1101/285981

**Authors:** Michaela Fenckova, Laura E.R. Blok, Lenke Asztalos, David P. Goodman, Pavel Cizek, Euginia L. Singgih, Jeffrey C. Glennon, Joanna IntHout, Christiane Zweier, Evan E. Eichler, Catherine R. von Reyn, Raphael A. Bernier, Zoltan Asztalos, Annette Schenck

## Abstract

**Background:** Although habituation is one of the most ancient and fundamental forms of learning, its regulators and relevance for human disease are poorly understood.

**Methods:** We manipulated the orthologs of 286 genes implicated in intellectual disability (ID) with or without comorbid autism spectrum disorder (ASD) specifically in *Drosophila* neurons, and tested these models in light-off jump habituation. We dissected neuronal substrates underlying the identified habituation deficits and integrated genotype-phenotype annotations, gene ontologies and interaction networks to determine the clinical features and molecular processes that are associated with habituation deficits.

**Results:** We identified more than 100 genes required for habituation learning. For the vast majority of these, 93 genes, a role in habituation learning was previously unknown. These genes characterize ID disorders with macrocephaly/overgrowth and comorbid ASD. Moreover, ASD individuals from the Simons Simplex Collection (SSC) carrying damaging *de novo* mutations in these genes exhibit increased aberrant behaviors associated with inappropriate, stereotypic speech. At the molecular level, ID genes required for normal habituation are enriched in synaptic function and converge on Ras-MAPK signaling. Both increased Ras-MAPK signaling in GABAergic and decreased Ras-MAPK signaling in cholinergic neurons specifically inhibit the adaptive habituation response.

**Conclusions:** Our work supports the relevance of habituation learning to autism, identifies an unprecedented number of novel habituation players, supports an emerging role for inhibitory neurons in habituation and reveals an opposing, circuit-level-based mechanism for Ras-MAPK signaling. This establishes habituation as a possible, widely applicable functional readout and target for pharmacologic intervention in ID/ASD.

## Introduction

Habituation is one of the most ancient and fundamental forms of learning, conserved across the animal kingdom (1). It causes an organism’s initial response to repeated meaningless stimuli to gradually decline. Learning to ignore irrelevant stimuli as a result of habituation is thought to represent a filter mechanism that prevents information overload, allowing for selective attention, thereby focusing cognitive resources on relevant matters. Habituation learning has been proposed to represent an important prerequisite for higher cognitive functions (2–4). In line with this, habituation in infants correlates better than other measures with later cognitive abilities (5). However, key players and molecular mechanisms underlying habituation are poorly understood (6).

In humans, deficits in habituation have been reported in a number of neuropsychiatric and behavioral disorders. In particular, defective cortical filtering of sensory stimuli and information overload, as expected to arise from habituation deficits, are thought to represent mechanisms contributing to autism spectrum disorder (ASD) (7, 8). A decreased ability to habituate has been described in a fraction of ASD individuals (9–11), but has not been connected yet to specific genetic defects, with a single exception. Recently, two independent studies demonstrated habituation deficits in patients with Fragile X syndrome, the most common monogenic cause of intellectual disability (ID) and ASD (12, 13), confirming previously reported habituation deficits in Fmr1 KO mice (14, 15). Habituation deficits have also been reported in a limited number of other ID or ASD (ID/ASD) disease models (16–19).

Because assessing human gene function in habituation is challenging, we utilized a cross-species approach. We apply light-off jump habituation in *Drosophila* to increase our knowledge on the genetic control of habituation and, at the same time, to address the relevance of decreased habituation in ID and in comorbid ASD disorders. Since ID is present in 70% of individuals with ASD (20), monogenic causes of ID provide a unique molecular windows to ASD pathology (21). *Drosophila* is a powerful, well-established model for ID (22–24) and offers genome-wide resources to study gene function in large scale (25, 26). Several forms of habituation have been established in *Drosophila* (27–31). Deficits in light-off jump habituation have already been reported in several ID models (23, 32–36) and in classical learning and memory mutants (28, 31). Moreover, this form of habituation can be assessed in a high-throughput manner. In the light-off jump paradigm, the initial jump response to repeated light-off stimuli gradually wanes, as has been demonstrated not due to sensory adaptation (a decrease in detecting the stimulus) or motor fatigue (a decrease in the ability to execute the response) but as a result of learned adaptation of the startle circuit (31). This behavior meets all habituation criteria (37), including spontaneous recovery and dishabituation with a novel stimulus (31, 38).

Here, we use inducible RNA interference (RNAi) in *Drosophila* to systematically assess the role of *Drosophila* orthologs of 286 genes that are well-established to cause ID in humans when mutated (hereinafter referred to as ID genes). 68 of them (20%) have also been implicated in ASD (39, 40) (**Table S1**), hereinafter referred to as ID plus ASD-associated genes.

## Methods and Materials

### Investigated ID genes

A systematic source of ID genes and their *Drosophila* orthologs is available online (SysID database, sysid.cmbi.umcn.nl (41)). We investigated the *Drosophila* orthologs of 286 human ID genes from the SysID category primary ID genes (**Table S1**) (containing mutations with robust published evidence for causality, see **Supplemental Methods (SM)**). SysID inclusion criteria and in/exclusion criteria of experimentally investigated genes are indicated in the **SM**). In brief, the vast majority of genes are from the first data freeze of the SysID database (status of mid 2010). Genes have been included based on conservation in *Drosophila*, available tools (RNAi) from large-scale resources and viability as a prerequisite for behavioral testing. No selection was performed.

### Light-off jump habituation assay

3- to 7-day-old flies were subjected to the light-off jump habituation paradigm in two independent 16-unit light-off jump systems (manufactured and distributed by Aktogen Ltd.). After 5 min adaption, flies were simultaneously exposed to a series of 100 light-off pulses (15 ms) with 1 s interval. The noise amplitude of wing vibration during jump responses was recorded. An appropriate threshold (0.8 V) was applied to filter out background noise. Data were collected by a custom-made Labview Software (National Instruments). Flies were considered as habituated when not jumping in five consecutive light-off trials (no-jump criterion). Habituation was quantified as the number of trials required to reach the no-jump criterion (Trials To Criterion (TTC)).

Information about the identification of *Drosophila* orthologs, proposed disease mechanism, *Drosophila* stocks, phenotype reproducibility, validation of the automated jump scoring and of jump specificity, fatigue assay, quality criteria for RNAi lines, annotation of ID plus ASD associated genes, enrichment analysis, comparison of behavior and cognition in ASD individuals from the SSC, molecular interaction network, clustering, physical interaction enrichment (PIE), data visualization and statistics are described in the **SM**.

## Results

### Systematic identification of habituation deficits in *Drosophila* models of ID

To identify novel genes implicated in habituation, we systematically investigated the role of 278 *Drosophila* orthologs representing 286 human ID genes in the light-off jump habituation paradigm. We induced neuron-specific knockdowns of each ID gene ortholog by RNAi (25) using 513 RNAi lines fulfilling previously established quality criteria (41, 42), with two independent constructs per gene whenever available. These were crossed to the panneuronal elav-Gal4 driver line (see **SM**). Knockdown is a suitable approach for modeling of the here-investigated human disease conditions since (partial) loss of function is considered to be the underlying mechanism in the vast majority of these disorders (42) (**Table S1**). Restricting gene knockdown to neurons eliminates potential effects on viability or behavioral performance originating from an essential role of genes in other tissues and establishes neuron-autonomous mechanisms.

Knockdown and control flies of identical genetic background were subjected to a series of 100 light-off stimuli, hereinafter referred to as trials, in the light-off jump habituation paradigm. The screening procedure and paradigm allowed us to distinguish the following parameters: viability, initial jump response (percentage of flies that jumped in at least one of the first five trials), and premature and reduced habituation, with the latter representing the learning-defective phenotype category of main interest. Genotypes with an initial jump response ≥50% but premature habituation were subjected to a secondary assay to exclude fatigue as a confounder of premature habituation (see **SM**, **Table S2** and **Figure S4**). Based on these parameters, genes were assigned to at least one of four phenotype categories (**Figure 1A**): (1) “not affected”: (both) tested RNAi lines targeting such genes were viable, showed good initial jump response, and had no significant effect on habituation (based on the FDR-corrected p-value (p_adj_), see **SM**); (2) “non-performers”: at least one RNAi line led to lethality, poor jump response (<50% initial jumpers), or premature habituation because of increased fatigue; (3) “habituation deficient”: at least one RNAi line showed good initial jump response but failed to suppress the response with the increasing number of light-off trials (based on p_adj_); and (4) “premature habituation”: at least one RNAi line showed good initial jump response followed by faster decline (based on p_adj_), without fatigue being detectable in the secondary assay. Still, this latter phenotype category can result from other defects than improved habituation, and will be investigated elsewhere. In this study we focus on habituation deficits (3), corresponding to the phenotype that has been shown in ID and ASD (9–13).

**Figure 1.**
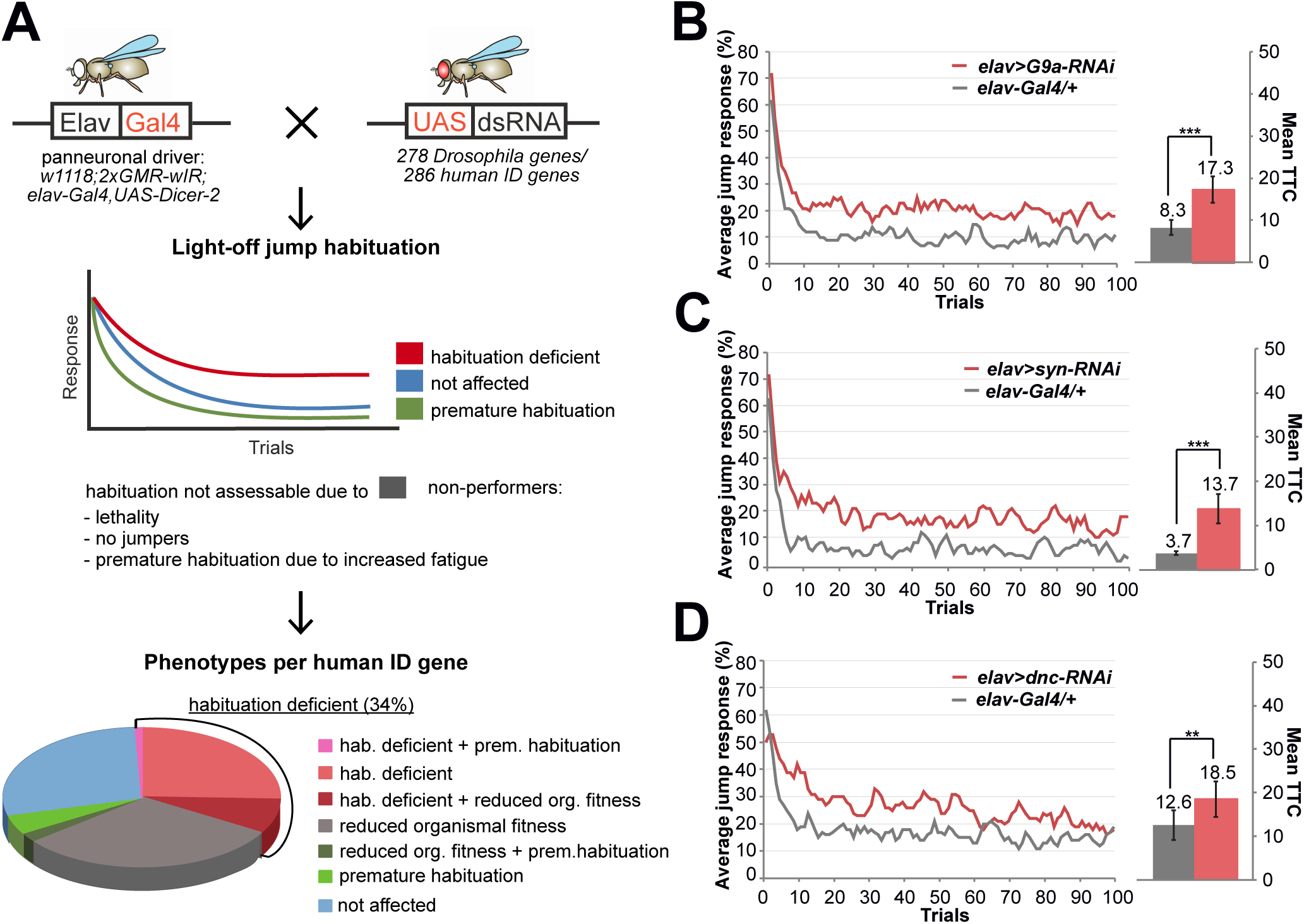
Habituation screen of intellectual disability genes, phenotype distribution and proof of principle. (A) Procedure, phenotype categories and phenotype distribution of the light-off jump habituation screen. Knockdowns that resulted in lethality, no jumper phenotype (defined as less than 50% flies jumping in at least one of the first five light-off trials) or premature habituation plus increased fatigue were assigned to the category “non-performers” and their habituation was not further analyzed. Other phenotype categories are “habituation deficient”, “not affected”, and “premature habituation” (the latter if no fatigue was detected in secondary assay, see example in **Figure S4**). *Drosophila* orthologs of 34% of the investigated human ID genes were associated with defects in habituation learning. See also **Table S2, S3**. (B, C, D) Defective habituation upon neuron-specific RNAi-mediated knockdown of *G9a, Synapsin (syn)*, and *dunce (dnc)* (*2xGMR-wIR/+; UAS-RNAi/elav-Gal4, UAS-Dicer-2*, in red) compared to their respective genetic background controls (*2xGMR-wIR/+; elav-Gal4, UAS-Dicer-2/+*, in gray). Jump response curves show the average jump response (% of jumping flies) over 100 light-off trials at 1 s inter-trial interval). Mean TTC: the mean number of trials that flies needed to reach the no-jump criterion (see **Methods and Materials**) presented as Mean TTC ± SEM. *** p_adj_<0.001, ** p_adj_<0.01, based on FDR-corrected lm analysis. A complete list of ID genes with previously identified habituation defects is provided as **Table S8**, adding further proof of principle.

We validated the experimental approach to identify genes which, if manipulated, cause habituation deficits (hereinafter referred to as habituation deficient genes) by recapitulating published habituation deficits of *Drosophila* ID null mutant models *G9a* (23) and *Synapsin* (43), and of the classical learning and memory mutant *dunce* (28, 44, 45) (**Figure 1B,C,D)**. This demonstrated that light-off jump habituation upon RNAi can efficiently identify genetic regulators of habituation learning. We also validated the technical accuracy of the automated jump scoring methodology by comparing automated and manually assessed jumping of controls and a number of ID models (**SM**, **Figure S1**).

In our screen, we found that the *Drosophila* orthologs of 98 human ID genes (35% of all investigated orthologs) are required, in neurons, for habituation learning. This phenotype represents a highly specific defect in behavioral adaptation to the stimulus; flies keep on jumping in response to the repetitive light-off stimulus, illustrating that they do not suffer from broad neuronal transmission deficits (which would disable jumping), fatigue, sensory or other deficiencies. No excessive locomotion was observed when handling the flies, and no stimulus hypersensitivity or random jumping was found (see **SM** and **Figure S2, S3** for validation of light-off jump habituation assay specificity). 27% of ID gene orthologs had no effect on habituation, 41% fell into the category of “non-performers”, and 8% showed “premature habituation” without detectable fatigue. The complete list of habituation screen results and distribution of human ID genes in phenotype categories can be found in **Table S2, S3**. The screen thus identified nearly a hundred orthologs of disease genes controlling habituation learning.

### Habituation deficits characterize ID genes with synaptic function

We first asked whether genes characterized by habituation deficits in *Drosophila* converge on specific biological process. ID genes are known to be enriched in a number of biological processes, but which are important for habituation? Performing an enrichment analysis of ID-enriched Gene Ontology-based (GO) categories (see **SM**) against the background of the investigated ID genes, we found that “habituation deficient” genes are significantly enriched in a sole GO-based category: processes related to the synapse (22/44 ID genes, E=1.59, p=0.024, **Figure 2**, **Table S4**). No enriched GO terms were found in the “not affected” category. Together, our results support synaptic processes to be crucial for habituation, as previously shown for other forms of this behavior (46, 47).

**Figure 2.**
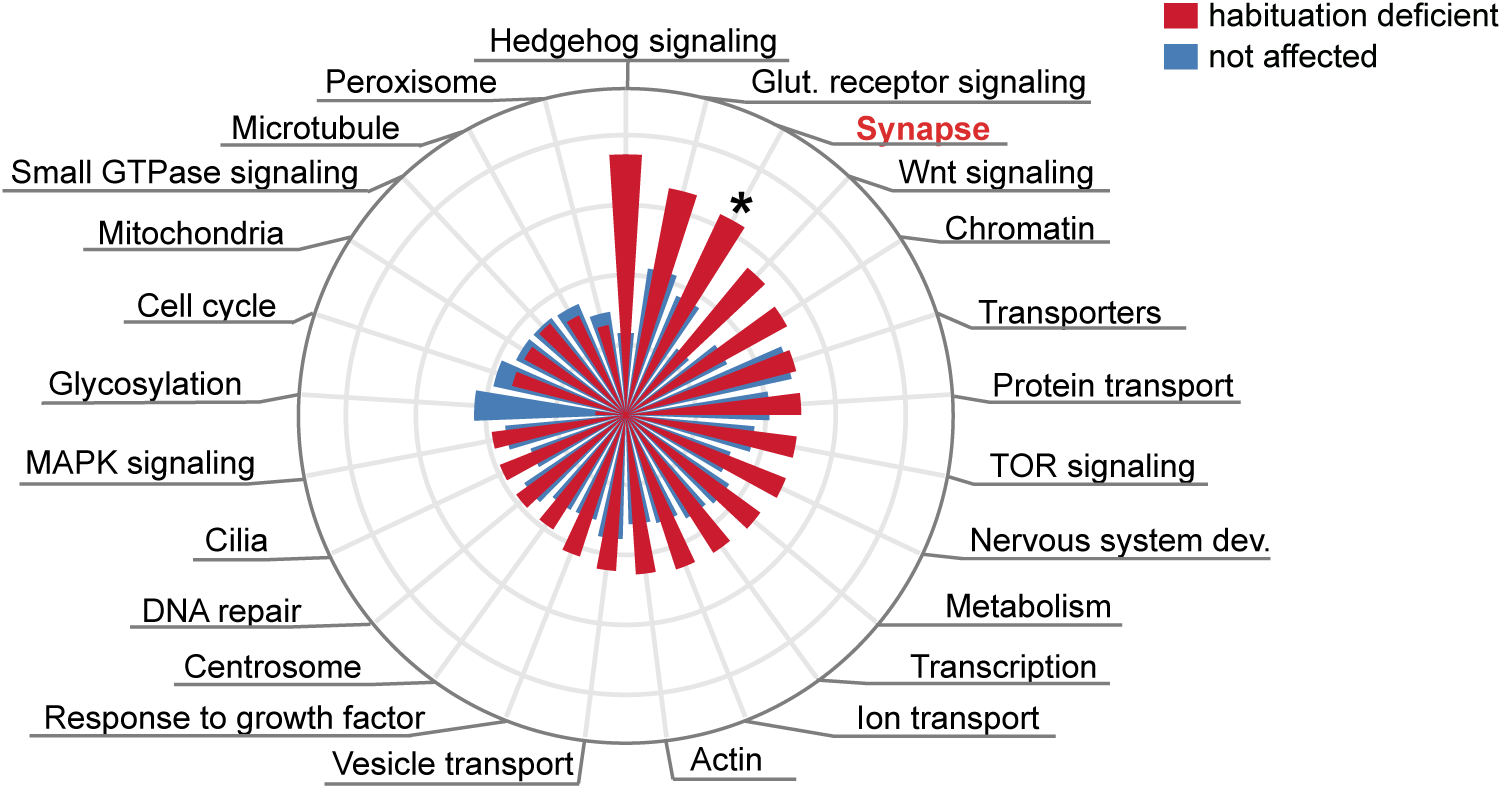
Habituation deficits in *Drosophila* characterize ID genes with synapse-related functions. Of 25 gene ontology (GO)-based processes, “habituation deficient” genes are specifically and significantly enriched in processes related to synapse (E=1.59, p=0.024). Genes with no effect on habituation do not show significant enrichment in any GO process. * p<0.05, based on Fisher’s exact test. All enrichment scores, p-values and enriched genes are listed in **Table S4**.

### *Drosophila* habituation deficits characterize ID genes associated with macrocephaly

To understand whether habituation deficits in *Drosophila* represent a proxy of specific phenotypes in human individuals, we performed enrichment analysis among ID-associated clinical features (41). We found that orthologs of ID genes characterized by habituation deficits in *Drosophila* are specifically enriched among ID genes associated with macrocephaly/overgrowth (**Figure 3**, E=2.19, p=0.018, **Table S4**). In contrast, ID genes characterized as “non-performers” show enrichment in different, severe ID-associated features such as endocrine, limb and eye anomalies, brain malformations and obesity (**Figure S5**, **Table S4**). Moreover, ID genes not giving rise to habituation deficits (“not affected” category) did not show any enrichment among ID-associated clinical features (**Figure 3**, **Table S4**).

**Figure 3.**
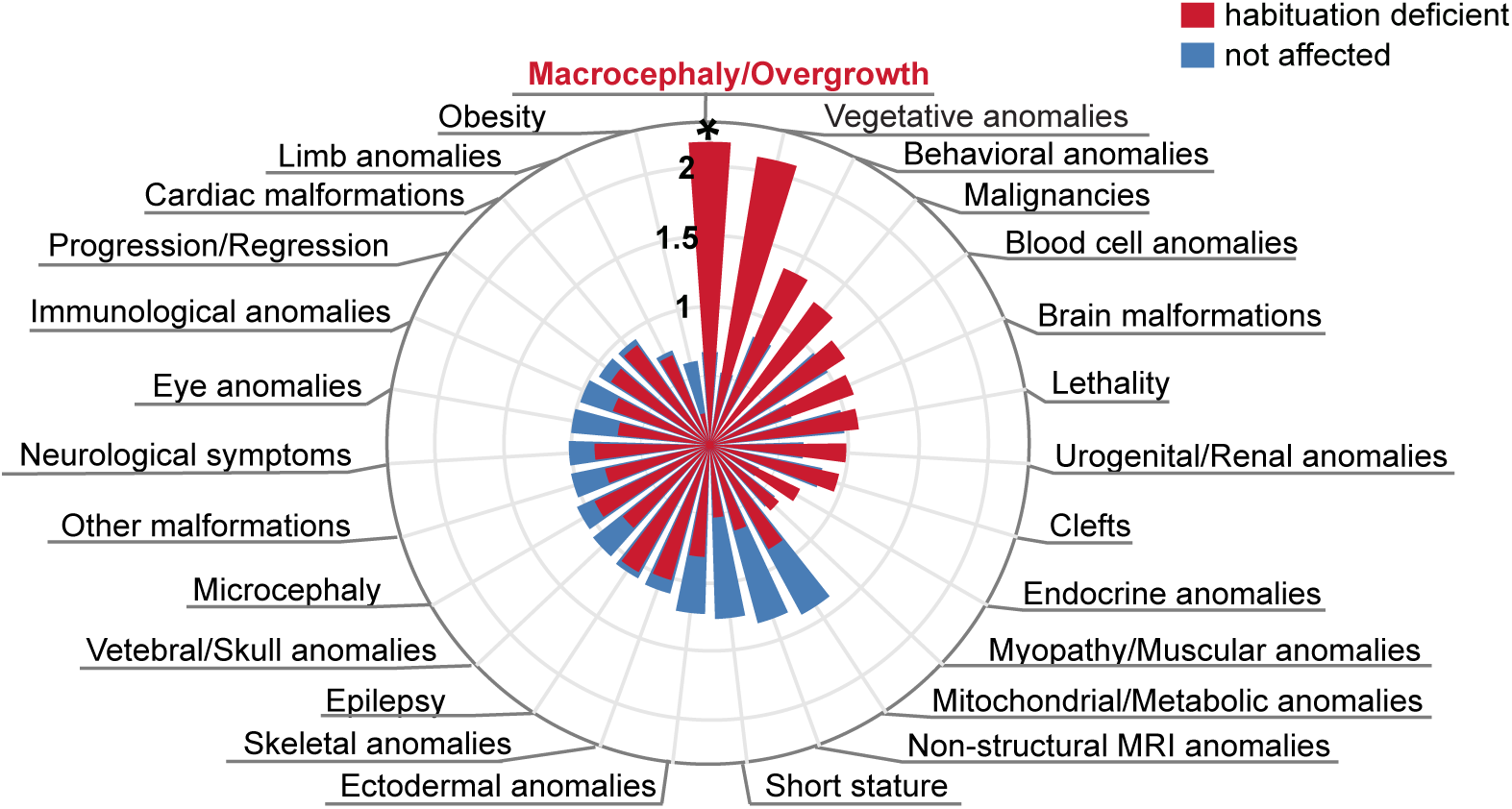
Habituation deficits in *Drosophila* characterize ID genes associated with macrocephaly in humans. Enrichment of *Drosophila* phenotype categories across 27 ID-accompanying clinical features (41). “Habituation deficient” genes show specificity for macrocephaly and/or overgrowth (E=2.19, p=0.018) ** p<0.01, * p<0.05, based on Fisher’s Exact test. For enrichment among the “non-performers” category, see **Figure S5**. Enrichment scores, p-values and enriched genes are listed in **Table S4**.

### Habituation deficits characterize ID genes associated with ASD and deficits in specific ASD-relevant behavioral domains

There is a long-known relationship between macrocephaly and autism (48). For this reason and because of the potential relevance of habituation deficits to ASD (9–11), we decided to further investigate the relationship of *Drosophila* habituation and human ASD. We used the Simons Simplex Collection (SSC) (40), a genetically and phenotypically well-characterized cohort of sporadic ASD individuals. We matched genes with likely gene-disrupting (LGD) and likely damaging *de novo* mutations (49, 50) in this ASD cohort to those included in our experimental *Drosophila* habituation approach. 47 ASD individuals carried mutations in 33 of the investigated genes (**Table S5**). We first asked whether these ID plus ASD-associated genes preferentially fall into a specific *Drosophila* phenotype category. They are significantly enriched among the genes that in *Drosophila* caused habituation deficits (**Figure 4A,** E=1.64, p=0.029, **Table S4**, ASD SSC). Independently, significant enrichment was obtained for high-confidence ID plus ASD-associated genes identified from the SFARI database (39) (38 investigated genes, **Figure 4B,** E=1.65, p=0.016, **Table S4**, ASD SFARI), suggesting a relationship between *Drosophila* habituation deficits and human ASD.

**Figure 4.**
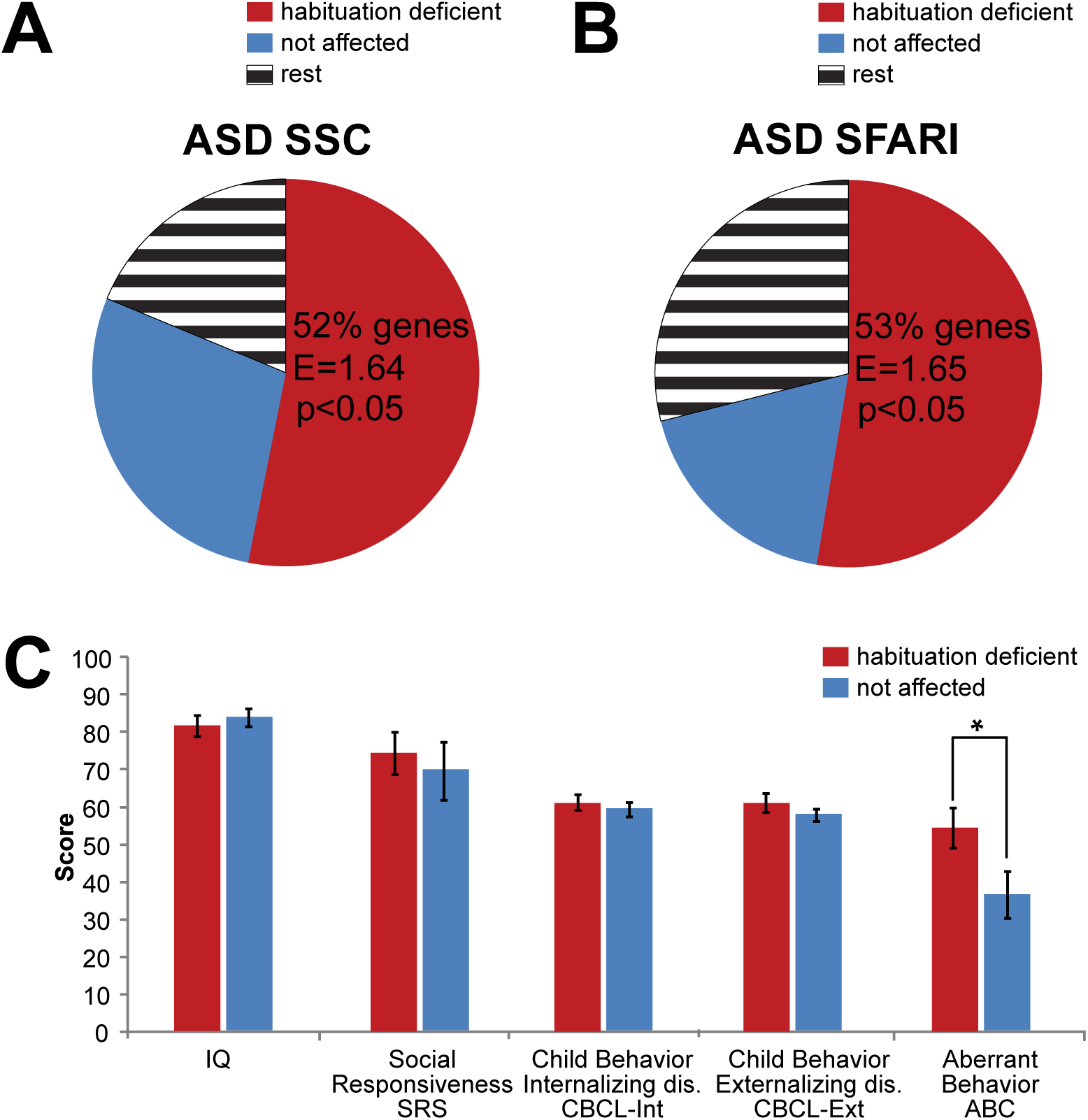
Habituation deficits in *Drosophila* characterize ID genes associated with ASD and deficits in specific behavioral domains. (A,B) Enrichment of *Drosophila* phenotype categories “habituation deficient” and “not affected” in ID plus ASD-associated genes identified in SFARI database (ASD SFARI, E=1.65, p=0.016, (A)) and SSC cohort (ASD SSC, E=1.64, p=0.029 (B)). Circles represent total number of tested ID plus ASD-associated genes. (C) Genes associated with “habituation deficient” versus “not affected” phenotype categories in *Drosophila* show tendency for more aberrant behaviors on the ABC (p=0.04) in the ASD SSC cohort. Data presented as mean score ± SEM. * p<0.05, based on MANOVA. See also **Table S5** (list of ASD SSC and ASD SFARI genes) and **Table S6** (complete MANOVA results).

To further characterize the relationship between *Drosophila* habituation and human phenotypes, we divided the SSC individuals into two distinct clusters based on their habituation phenotype in the corresponding fly models: habituation deficits (N=22 individuals, 17 genes) and no habituation deficits (N=12 individuals, 9 genes) (**Table S5;** another N=13 individuals, 7 genes fall into the non-informative phenotype groups “non-performers”/“premature habituation”). We compared both groups across five broad quantitative measures of behavior and cognition: cognitive ability (full-scale IQ); Social Responsiveness Scale (SRS); depression and anxiety (Child Behavior Checklist Internalizing Disorders, CBCL-Int); impulsivity, attention and conduct (Child Behavior Checklist Externalizing Disorders, CBCL-Ext); and atypical behavior (Aberrant Behavior Checklist, ABC). There was no significant difference for IQ (p=0.61), SRS (p=0.62), CBCL-Int (p=0.59) or CBCL-Ext (p=0.37), but a trend for ABC (p=0.04; **Figure 4C**, **Table S6)**. This effect is mainly driven by the ABC subdomain of inappropriate, stereotypic speech (p=0.0003), not from the subdomains of irritability (p=0.1), hyperactivity (p=0.86), lethargy (p=0.54) or stereotypy (p=0.91) (**Table S6**). In summary, these data indicate that habituation deficits in *Drosophila* are relevant to ASD-implicated genes. They also suggest that SSC individuals carrying *de novo* mutations in genes associated with habituation deficits in *Drosophila* show a higher rate and/or severity of atypical behaviors associated with inappropriate and stereotypic speech.

### Molecular networks and modules underlying habituation

With the rich repertoire of nearly a hundred genes required for habituation that moreover show specificity for ASD and synapse function, we set out to determine the molecular pathways these genes are operating in. ID gene products are significantly interconnected via protein-protein interactions (51, 52). Consistent with previously published findings (41), ID genes investigated in our screen are 1.69 times enriched in interactions compared to 1000 randomly chosen protein sets of the same size and number of known interactions (physical interaction enrichment (PIE) score (53) =1.69; p<0.001). To identify biologically relevant modules, we resolved this network into communities with even tighter interconnectivity using unsupervised community clustering (54). This analysis resulted in 26 communities containing 109 proteins (**Figure 5A, Table S7**). Their proximity and specificity for ID-enriched GO-based processes are depicted in **Figure S6**. Mapping “habituation deficient” genes onto the communities (**Figure 5A, red circles**) highlighted modules with high incidence of habituation deficits (**Figure 5A**).

**Figure 5.**
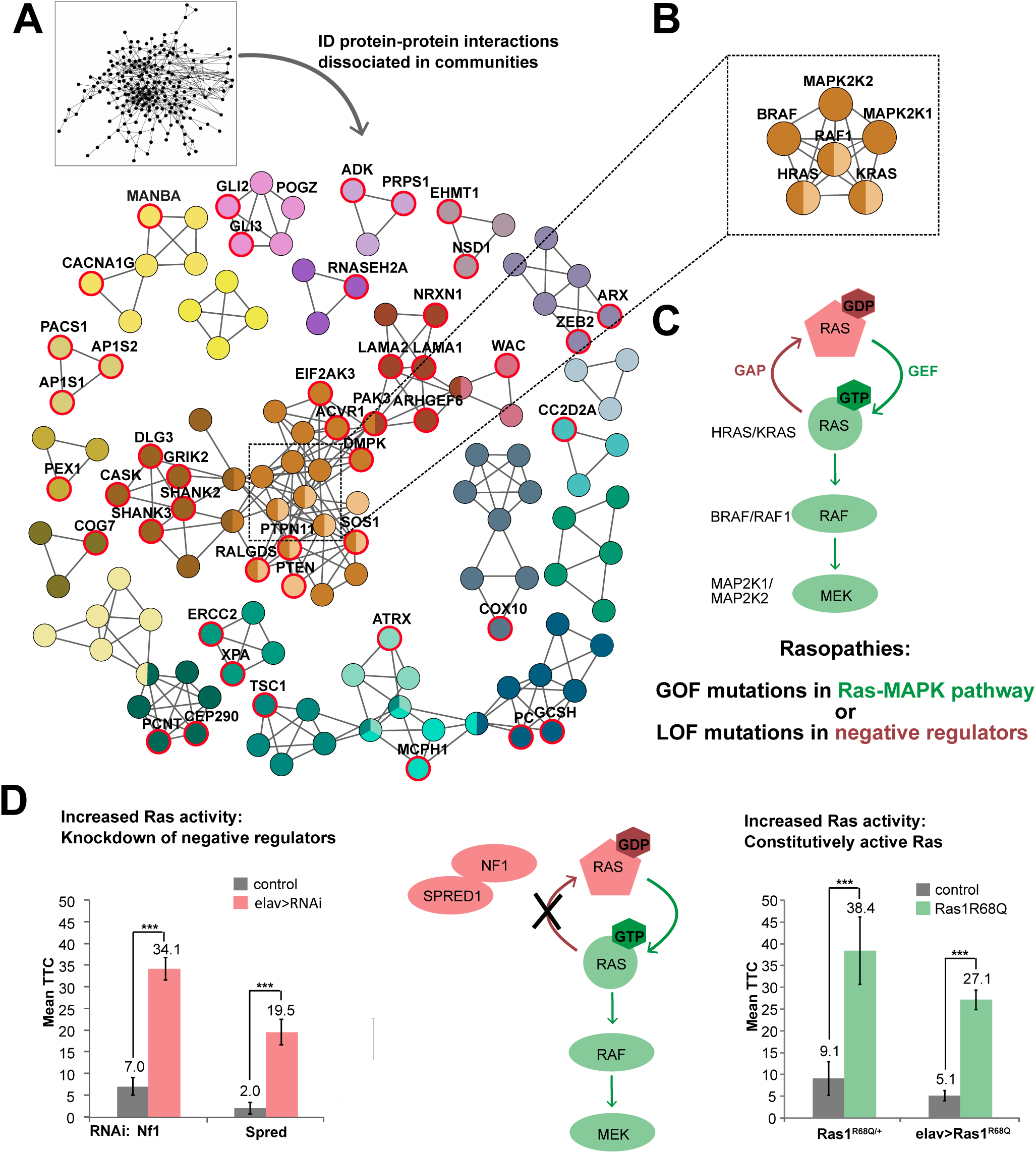
A central role for Ras-MAPK signaling in habituation learning. (A) Highly connected communities identified by unbiased community clustering, colored by their functional proximity (**Figure S6**). Red circles and gene names highlight nodes representing “habituation deficient” genes. For complete list of communities and genes see **Table S7**. (B) Nodes connecting four communities from the central module represent the core components of Ras-MAPK signaling. (C) Schematic representation of Ras-MAPK signaling and associated mechanisms in ID disorders called ‘Rasopathies’. (D) Increasing Ras signaling by inducing either loss of function of negative Ras regulators (left side of pathway scheme) or by constitutively activating Ras (right side) disrupts habituation learning. Left: Defective habituation upon neuron-specific knockdown of negative Ras regulators, *Nf1* (*2xGMR-wIR/+; Nf1-RNAi*^*vdrc35877*^*/elav-Gal4, UAS-Dicer-2*, N=72, in red) and *Spred* (*2xGMR-wIR/+; Spred-RNAi*^*vdrc18024*^*/elav-Gal4, UAS-Dicer-2*, N=73, in red), compared to their corresponding genetic background controls (*2xGMR-wIR/+; elav-Gal4, UAS-Dicer-2/+*. N: 55, 20, in gray). *** p_adj_<0.001, based on lm analysis and FDR correction in the screen (see **Methods and Materials**). Right: Defects in habituation learning in a heterozygous, constitutively active *Ras* mutant (*Ras1*^*R68Q/+*^, N=55, in green) compared to its genetic background control (N=43 in gray), and upon neuron-specific expression of *Ras1*^*R68Q*^ (*elav>Ras1*^*R68Q*^: *UAS-Ras1*^*R68Q*^*/2xGMR-wIR; elav-Gal4, UAS-Dicer-2/+*, N=52, in green) compared to its genetic background control (*2xGMR-wIR/+; elav-Gal4, UAS-Dicer-2/+*, N=34, in gray). *** p<0.001, based on lm analysis. Data presented as Mean TTC ± SEM.

### A key role for ID and ASD-associated Ras signaling in habituation

Five communities form a large, interconnected module with high incidences of habituation deficits. However, the tightly interconnected hub at its center is characterized by the absence of habituation deficits (**Figure 5A, square)**. This hub represents the key proteins of Ras-MAPK signaling (**Figure 5B**). This pathway, best known for its role in cancer, underlies a group of disorders collectively referred as Rasopathies. Importantly, while 92% of the modeled ID disorders are thought to result from loss of function of the underlying genes, Rasopathies are caused by gain-of-function mutations in the core pathway (**Figure 5C, Table S1**). The utilized RNAi approach, despite addressing gene function, did thus not recapitulate the molecular pathology of these specific cognitive disorders. However, Rasopathies can also result from loss of function in negative regulators of the pathway. We therefore asked whether the same genetic mechanisms that cause Rasopathies in humans also hold true for habituation deficits in *Drosophila*. In our screen, we tested habituation of two negative regulators of Ras: NF1 (*Drosophila* Nf1) (55) and SPRED1 (*Drosophila* Spred) (56, 57). Panneuronal knockdown of either regulator caused strong habituation deficits (**Figure 5D, in red**). We therefore tested a constitutively active *Ras* mutant, *Ras1*^*R68Q*^ (58). Heterozygous *Ras1*^*R68Q*^ flies showed strong habituation deficits compared to the control flies with the same genetic background (p=3.56×10^-9^; **Figure 5D, in green**). The same was true when we overexpressed, specifically in neurons, *Ras1*^*R68Q*^ allele from an inducible transgene (p=1.96×10^-6^; **Figure 5D, in green**). We conclude that increased activity of Ras, causing Rasopathies and associated cognitive deficits in humans, causes habituation deficits in *Drosophila*.

### Habituation-inhibiting function of increased Ras-MAPK signaling maps to inhibitory/GABAergic neurons

We next aimed to identify in which type of neurons the habituation-inhibiting function of Ras-MAPK signaling resides. Because the well-characterized neurons of the giant fiber circuit controlling the light-off jump response are cholinergic (59), just as the majority of excitatory neurons in *Drosophila*, we first tested whether increased Ras-MAPK signaling activity would induce habituation deficits when directed to cholinergic neurons. For this, we adopted the knockdown of negative Ras regulators (*Nf1, Spred*), expressed constitutively active *Ras1* (*Ras1*^*R68Q*^), and tested expression of a gain-of-function allele of *Raf* (*Raf*^*GOF*^), a downstream mediator of Ras signaling. None of these, when driven by the cholinergic Cha-Gal4 driver, recapitulated the panneuronally evoked habituation deficits (**Figure 6A**).

**Figure 6.**
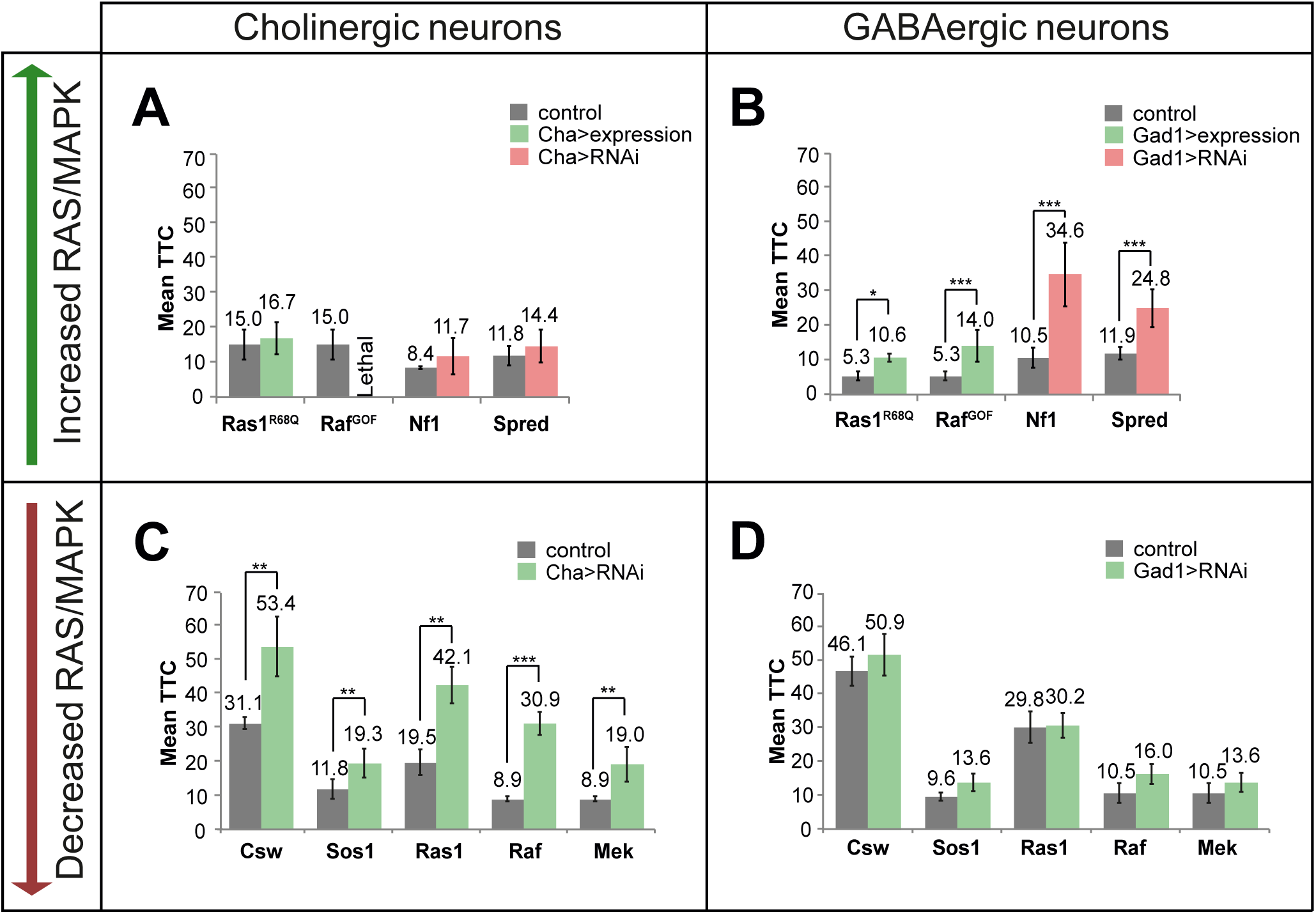
Dual, opposing role of Ras-MAPK signaling in GABAergic and cholinergic neurons in the regulation of habituation learning. (A) No effect on habituation of *Ras1*^*R68Q*^ (N=51, in green), *Nf1-RNAi* (N=38, in red), and *Spred-RNAi* (N=55, in red) upon expression in cholinergic neurons compared to their respective genetic background controls (*Cha-Gal4/+*; *2xGMR-wIR/+*, N: 54, 45, 54 in gray). Expression of *Raf*^*GOF*^ in cholinergic neurons resulted in lethality. (B) Defective habituation of *Ras1*^*R68Q*^ (N=52, in green), *Raf*^*GOF*^ (N=57, in green), *Nf1-RNAi* (N=55, in red), and *Spred-RNAi* (N=37, in red) on habituation upon expression in GABAergic neurons compared to their respective genetic background controls (*Gad1-Gal4/+*; *2xGMR-wIR/+*, N: 50, 50, 39, 58 in gray). (C) Defective habituation of *Csw-RNAi* (*UAS-Csw-RNAi*^*vdrc21756*^*/Y; Cha-Gal4/+*; *2xGMR-wIR/+*, N=58), *Sos1-RNAi* (*UAS-Sos1-RNAi*^*vdrc42848*^*/Cha-Gal4; 2xGMR-wIR/+*, N=56), *Ras1-RNAi* (*UAS-Ras1-RNAi*^*vdrc106642*^*/Cha-Gal4; 2xGMR-wIR/+*, N=55), *Raf-RNAi* (*UAS-Raf-RNAi*^*vdrc20909*^*/Cha-Gal4; 2xGMR-wIR/+*, N=59) and *Mek-RNAi* (*Cha-Gal4/+; UAS-Mek-RNAi*^*vdrc40026*^*/2xGMR-wIR*, N=58) in cholinergic neurons (in green) compared to their respective genetic background controls (*Cha-Gal4/+*; *2xGMR-wIR/+*, N: 62, 54, 34, 46, 46, in gray). (D) No effect on habituation of *Csw-RNAi* (N=58), *Sos1-RNAi* (N=51), *Ras1-RNAi* (N=53), *Raf-RNAi* (N=52) and *Mek-RNAi* (N=54) in GABAergic neurons (in green) compared to their respective genetic background controls (*Gad1-Gal4/+*; *2xGMR-wIR/+*, N: 60, 46, 54, 39, 39, in gray). Data presented as Mean TTC ± SEM. *** p<0.001, ** p<0.01, * p<0.05, based on lm analysis.

Because of the recently established role of GABAergic neurons in *Drosophila* olfactory and proboscis extension reflex habituation (29, 60, 61) and the emerging importance of GABA inhibition in autism (62), we next targeted GABA neurons using the Gad1-Gal4 driver and the same toolbox. This consistently induced habituation deficits in all tested conditions (**Figure 6B)**. We conclude that the habituation-inhibiting function of increased Ras-MAPK signaling maps to GABAergic neurons.

### Ras-MAPK signaling in cholinergic neurons is essential for habituation learning

Impaired jump response/increased fatigue associated with *Ras, Raf* and *Mek* knockdown in the screen could potentially mask an essential role for Ras signaling in habituation, in addition to the habituation-inhibiting function of increased Ras-MAPK signaling. In fact, our screen also identified habituation deficits upon RNAi of the positive Ras-MAPK regulators *Sos* and *Csw.* We therefore downregulated Ras-MAPK activity by crossing the UAS-based RNAi lines targeting *Sos* and *Csw*, but also RNAi lines targeting *Ras, Raf* and *Mek,* to the GABAergic driver Gad1-Gal4. We did not observe any detrimental effect on habituation (**Figure 6D**). In contrast, downregulating Ras-MAPK signaling in cholinergic neurons consistently prevented normal habituation learning (**Figure 6C**). We conclude that Ras-MAPK signaling is essential in cholinergic but not in GABAergic neurons. Thus, Ras-MAPK signaling plays a dual, opposing role in inhibitory versus excitatory neurons in habituation learning.

## Discussion

### *Drosophila* screen demonstrates that genes implicated in ASD are important for habituation learning

To systematically address the genetic basis of habituation deficits associated with neurodevelopmental disorders, we investigated 286 ID genes with a clear *Drosophila* ortholog in light-off jump habituation. Panneuronal knockdown of the orthologs of 98 ID genes specifically suppressed the adaptive habituation response to repeated stimulation without affecting organismal health or jump ability. Follow-up work on the Ras-MAPK pathway raised this number to 104. 93 of these are novel regulators of habituation, substantially exceeding the sum of previously known regulators of habituation across species and paradigms. Stringent criteria for RNAi specificity and correction for multiple testing (see **SM**) in our experiments ensured a minimal level of potential false positive discoveries. Of thirteen previously identified ID genes with habituation deficits, our screen confirmed ten (**Table S8**). Our approach and data, although based on experiments in another species, suggest that deficits in habituation learning are a widely affected mechanism in ID. Habituation deficits might be a hallmark of even more ID genes than determined here. In particular, the phenotype category of “non-performers” is likely to contain genes with promiscuous functions masking a specific role in habituation learning.

Enrichment analysis of ID-associated clinical features revealed that “habituation deficient” ID genes are preferentially characterized by macrocephaly/overgrowth, associated for long with ASD (48). Strikingly, we found that mutations in genes associated with *Drosophila* habituation deficits are significantly overrepresented among ID genes that are also implicated in ASD (52% (SSC cohort); 53% (SFARI database)). In comparison the frequency of habituation deficits among ID genes not associated with ASD is 24%. SSC individuals carrying mutations in these genes show a high rate and/or severity of aberrant behaviors associated with stereotypic speech. Habituation deficits thus represent a common phenotypic signature of ASD in *Drosophila* and highlight specific behavioral subdomains affected in ASD. Future work has to establish whether habituation deficits are a direct basis for these clinical features, or are one of many factors involved.

### Synapse-related processes and Ras-MAPK signaling play a key role in habituation

Synapse biology has been proposed to play a central role in ASD (63). Our data show that among the investigated disease genes, “habituation deficient” genes are specifically enriched in genes with synaptic function. This is in line with habituation representing a measurable form of synaptic plasticity (7, 47, 64).

Analyzing the distribution of “habituation deficient” genes in ID-specific molecular interaction networks, we discovered that they accumulate in a multiple-community module and connect to the Ras-MAPK pathway core proteins Ras, Raf and Mek (**Figure 5A,B**). We observed habituation deficits upon panneuronal knockdown of Ras negative regulators and panneuronal expression of the constitutively active *Ras* allele *Ras1*^*R68Q*^ (**Figure 5C**), demonstrating that increased Ras-mediated signaling causes habituation deficits. Moreover, proteins encoded by “habituation deficient” genes form a significantly interconnected module (**Figure 7**). The coherence of this module further supports the validity of the chosen RNAi approach to identify genes and molecular processes regulating habituation learning. The module contains a number of synaptic proteins (**Figure 7**) with not yet investigated roles in Ras signaling. It would be interesting to determine whether some of these enlarge the spectrum of diseases caused by deregulated Ras signaling.

**Figure 7.**
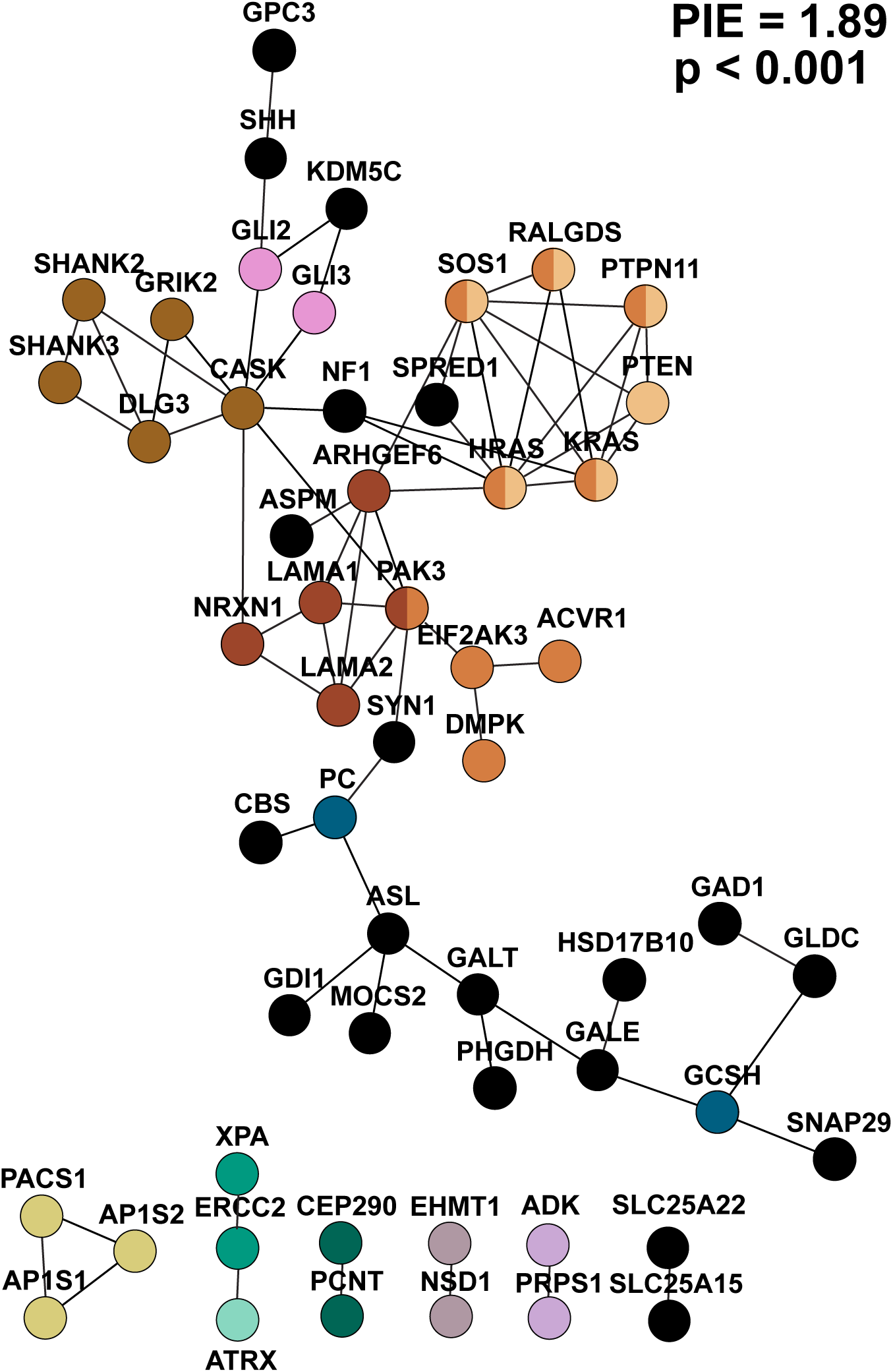
Connections between “habituation deficient” genes. Connections between “habituation deficient” genes, including *Ras*, identified in the reference network used for community clustering (See **SM**) with significantly increased connectivity (PIE score=1.89, p<0.001). Nodes are colored based on the community to which they belong. Nodes that represent “habituation deficient” genes but are not members of a community are labeled in black.

### Ras-MAPK signaling exerts a dual but opposing role in inhibitory versus excitatory neurons, a novel systems-level mechanism

Identification of neuronal substrates in which specific ID genes are required to warrant habituation learning is important fundamental problem. Restoring the function of affected neurons might also represent a suitable treatment strategy. The light-off jump startle circuit of *Drosophila* is relatively simple and its cholinergic nature is well described (59). However, it is not known how habituation of this circuit is regulated. The commonly accepted view regards synaptic depression in excitatory neurons, induced by repetitive stimulation, as the underlying mechanism (46, 65). This has recently been challenged by Ramaswami and colleagues who showed that plasticity of inhibitory, GABAergic neurons drives two non-startle types of habituation (60, 61). We found that increased activity of our identified key pathway, Ras-MAPK, in GABAergic but not in cholinergic neurons causes deficits in light-off jump habituation. Our results thus support inhibitory circuits as crucial components of habituation learning across different paradigms and sensory modalities. Further experiments are needed to establish the direct involvement of GABAergic signaling. At the same time, we identified that also decreased Ras-MAPK signaling activity can lead to habituation deficits. Yet, the neuronal substrates of these deficits are different and map to excitatory, cholinergic neurons. Although our experiments do not distinguish between developmental effects and acute circuit plasticity, the opposing role for Ras-MAPK signaling on habituation may provide new insights into mechanisms of neural plasticity in health and disease. It may also have crucial implications for treatment of Rasopathies. Future clinical trials, as opposed to those that broadly decreased Ras activity and failed (66), may need more attention towards restoring circuit function and balance.

### Translational value and application of cross-species habituation measures for diagnosis and treatment of ID and ASD

Based on our findings that habituation is widely affected in *Drosophila* models of ID, and that habituation deficits are particularly common among genes also implicated in ASD, we propose that disrupted habituation may be one of the mechanisms that contribute to ID/ASD pathology.

The emerging importance of inhibitory inputs for habituation ((29, 60) and this study) and sensory information filtering in the cortical centers of the brain (67, 68) suggests the existence of an overarching circuit-based mechanism responsible for prevention of inappropriate behavioral responses (7). Though our findings that habituation deficits in *Drosophila* correlate with increased rate and/or severity of atypical ASD-related behaviors compared to ID genes without habituation deficits should be replicated, we speculate that disrupted habituation arising from GABAergic defects may contribute to these ASD features. If future work can establish a substantial contribution of deficits in habituation learning to patient outcomes, cross-species habituation could become an attractive mechanism-specific functional readout—a pressing need for efficient personalized (pharmacological) treatment in the field of neurodevelopmental disorders. Implementing suitable low-burden protocols for habituation measures in clinical research and diagnostics of ID/ASD, such as those developed for investigation of habituation deficits in Fragile X syndrome (12, 13), will help to further delineate the affected cognitive domains that may correlate with or arise from deficient habituation. In future clinical trials, these could serve as objective and quantitative readouts for patient stratification in mechanism-based treatment strategies and for monitoring of drug efficacy. Dissection of the underlying defective mechanisms in *Drosophila* can at the same time identify novel targets for treatment, with high-throughput light-off jump habituation serving as a translational pipeline for drug testing.

## Acknowledgements

We thank Dr Erika Virágh and Enikő Csapó (Biological Research Centre, Szeged, Hungary), moreover Dr Judit Bíró and Márk Péter-Szabó (Voalaz Ltd., Szeged, Hungary) for their contribution to the validation of the *Drosophila* semi-automated light-off jump reflex habituation paradigm. We acknowledge the Vienna *Drosophila* Resource Center and Bloomington *Drosophila* Stock Center (NIH P40OD018537) for providing *Drosophila* strains. We thank the anonymous expert referees for constructive feedback. This research was supported in part by the European Union’s FP7 TACTICS, OPTIMISTIC, Aggressotype and MATRICS (HEALTH grant agreement numbers n°278948, n°305697, n°602805 and n°603016) to J.C.G., by a grant from the National Science Foundation (CBET-1747506) to C.R.v.R., by the FP7 large-scale integrated network Gencodys (HEALTH-241995) to Z.A. and A.S., by a TOP grant (912-12-109) from The Netherlands Organization for Scientific Research (NWO), by a Horizon 2020 Marie Sklodowska-Curie European Training Network grant (MiND, 643051), by a grant from the Jérôme Léjeune foundation, and by a grant awarded under the Australian National Health & Medical Research Council (NHMRC) Centre for Research Excellence Scheme (APP1117394) to A.S., and by U.S. National Institute for Mental Health (NIMH) funding (R01MH101221 to E.E.E. & R01MH100047 to R.B.). E.E.E is an investigator of the Howard Hughes Medical Institute. We are grateful to all of the families at the participating Simons Simplex Collection (SSC) sites, as well as the principal investigators (A. Beaudet, R. Bernier, J. Constantino, E. Cook, E. Fombonne, D. Geschwind, R. Goin-Kochel, E. Hanson, D. Grice, A. Klin, D. Ledbetter, C. Lord, C. Martin, D. Martin, R. Maxim, J. Miles, O. Ousley, K. Pelphrey, B. Peterson, J. Piggot, C. Saulnier, M. State, W. Stone, J. Sutcliffe, C. Walsh, Z. Warren, E. Wijsman). We appreciate obtaining access to phenotypic data on SFARI Base. Approved researchers can obtain the SSC population dataset described in this study (http://sfari.org/resources/simons-simplex-collection) by applying at https://base.sfari.org.

