## Supplementary material for "A hundred genes implicated in intellectual disability and autism regulate habituation learning and reveal an opposing role for Ras-MAPK signaling in inhibitory and excitatory neurons": Document S1

**Supplemental information**

**Videos:**

**Video S1**

Provided as a separate e-component file. Combined, synchronized video of an example null response of a control fly (*2xGMR-wIR/+; elav-Gal4, UAS-Dicer-2/+,* left)) and an example escape response of a knockdown fly (elav-Gal4/vdrc36328, right) to the light-off stimulus. Escape responses consist of a rapid middle leg extension (take off jump) and wing fluttering, illustrating the wing engagement detected by the automated light-off jump habituation system. The light-off stimulus is initiated at time 0.1 s as indicated by the disappearance of the green square in the lower right corner of the video (frame 300-301).

**Video S2**

Provided as a separate e-component file. Representative example of a simultaneous audio and video recording of a single control fly in the light-off jump habituation system, for detailed view. The fly housed in chamber 10 (plot 10) is exposed to 10 light-off stimuli with a 5 s interval between the stimuli. When the light-off stimulus is followed by a jump response, the noise threshold reaches the value of 0.8 V or higher (jump threshold) on the vertical axis and the graphical indicator next to plot 15 turns green, indicating the detection of a jump. After stimuli without a jump response, the noise amplitude remains below the jump threshold and no jump is detected. Note the slight delay between the video-taped jump and the graphical representation of the jump amplitude. The amplitude is being recorded for 500 ms after stimulus onset and is depicted on screen only after the 500 ms recording has been completed.

**Video S3**

Provided as a separate e-component file. Representative example of a simultaneous audio and video recording of an 8-vial unit of the light-off jump habituation system, as used to generate the data for quantitative comparison of manual (video) and automated (audio) jump scoring (**Figure S1**). 8 flies are housed in chambers represented by plots 9-16. To facilitate visibility of short light-off pulses, the light-off pulses were adapted to 40 ms in this movie (compared to 15 ms in **Video S2**, in the screen and in the validation experiments in **Figure S1**). For scoring of the videos, the video recording of the chambers was expanded to the full computer screen. Jumps and TTCs for each individual fly were manually scored, blinded to genotype and audio scores.

**Figures:**

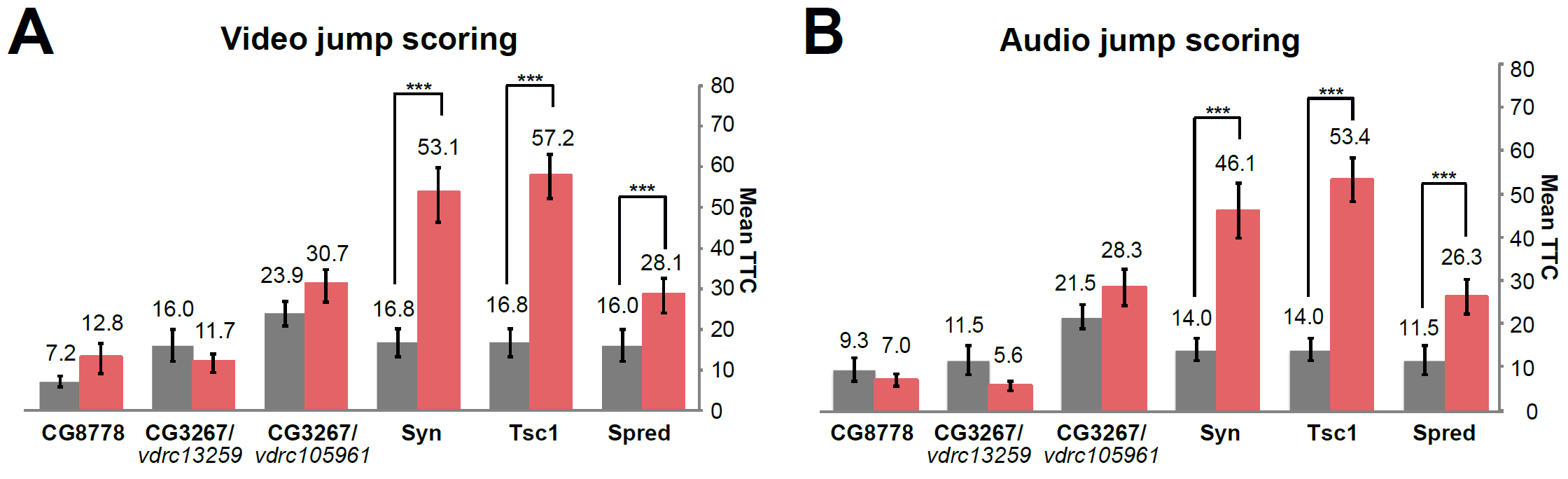

**Figure S1. Validation of habituation phenotypes obtained with the automated jump scoring**

Quantitative comparison of mean TTC values between controls (in gray) and knockdown flies (in red) in the same experiments, simultaneously recorded by video and audio. Three habituation deficient and three not affected genotypes were selected from the screen. (A) Video scoring of jump responses. No effect on habituation of *CG8778* (N=49, p=0.613), *CG3267^vdrc13259^* (N=48, p=0.735), *CG3267^vdrc105961^* (N=53, p=0.276) and habituation deficit of *Syn* (N=38, p=7.62x10^-5^), *Tsc1* (N=58, p=4.01x10^-7^), *Spred* (N=52, p=3.54x10^-4^) compared to the genetic background controls (*2xGMR-wIR/+; elav-Gal4, UAS-Dicer2/+*, N: 49, 51, 57, 50, 50, 51). (B) Audio scoring of jump responses. No effect on habituation of *CG8778* (N=46, p=0.844), *CG3267* (N=32, p=0.327), *CG3267* (N=49, p=0.497) and habituation deficit of *Syn* (N=38, p=7.62x10^-5^), *Tsc1* (N=58, p=4.01x10^-7^), *Spred* (N=52, p=3.54x10^-4^). compared to the genetic background controls (*2xGMR-wIR/+; elav-Gal4, UAS-Dicer2/+*, N: 46, 43, 53, 44, 44, 43). Data from two independent repeats are presented as Mean TTC ± SEM. *** p<0.001 based on lm analysis. These experiments illustrate the accuracy of the automated jump scoring and the reproducibility of results obtained in the screen.

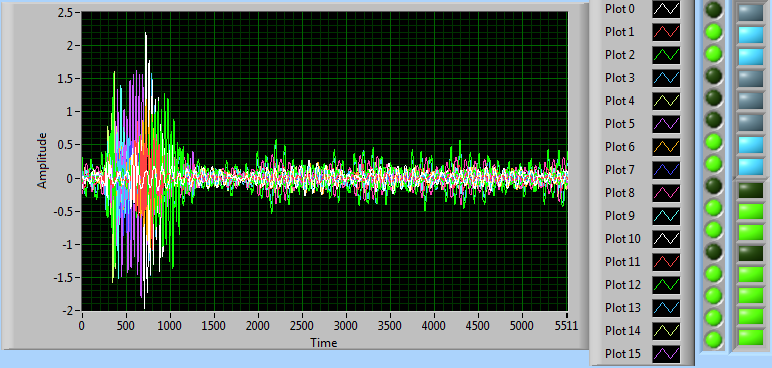

**Figure S2. Sound profile view of individual chambers during light-off jump habituation**

Overlay of individual sound/jumping profiles from the 16 chambers of the light-off jump system (color-coded), as visible to the experimenter during the experiment. Vertical axis shows the noise amplitude (in volts) as detected by two microphones per chamber (one plotted as positive amplitude, the other as negative). The horizontal axis shows the time after onset of the stimulus (1 unit corresponds to ~1/10 ms). Chambers in which the noise amplitude crosses the “jump” threshold of 0.8 V are labeled in blue or green (on the right). Chambers in this setup are isolated from each other, assuring that the recorded sound is chamber-specific and that flies do not visually react to each other. Note that jump responses in the individual chambers) occur fast and synchronized. Random jumping was not observed for any of the tested genotypes including genetic background controls (see also **Figure S3A**).

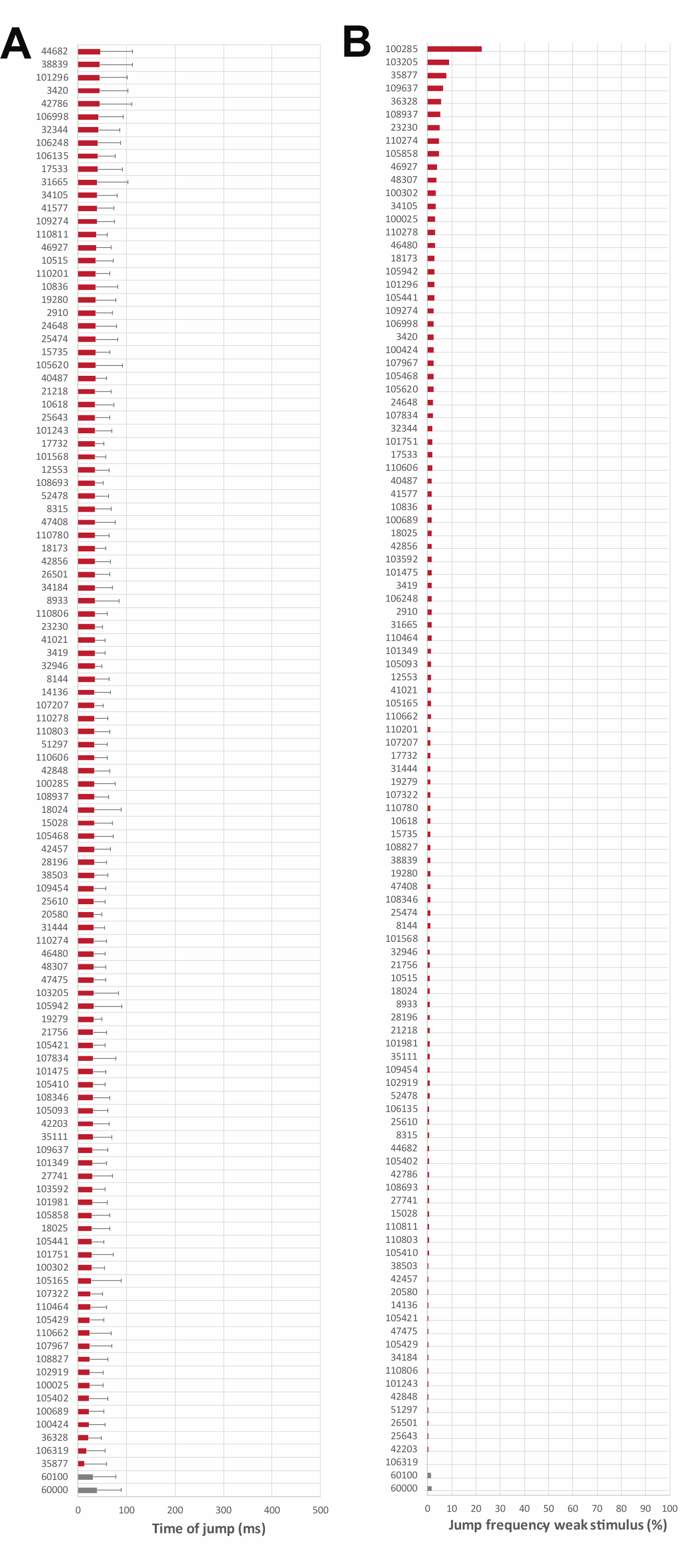
**Figure S3. Specificity of jumps and sensitivity to the light-off stimulus**

(A) Mean time of jump after light-off stimulus (horizontal, in ms) was calculated for the two controls (bottom, in grey) and all “habituation deficient” genotypes (in red) to exclude random jumping. All genotypes jumped within the first 100 ms after the light-off stimulus, demonstrating that jumps do occur in immediate response to stimulation; random jumping did not occur in any genotype. Genotypes are labeled with the vdrc order numbers of the respective RNAi line. Error bars represent standard deviation. (B) Jump frequency in response to a weaker stimulus. The light intensity in these experiments is decreased/dimmed to 70%, a stimulus intensity just below the threshold of eliciting a jump response. A slightly increased sensitivity would thus, result in jumping. All tested flies were exposed to a series of 10 weak stimuli with a 5 s interval between stimuli (preventing habituation, as in the fatigue assay). Controls (bottom, in grey) and all “habituation deficient” genotypes (in red) are shown, their ump frequency in % is plotted horizontally.

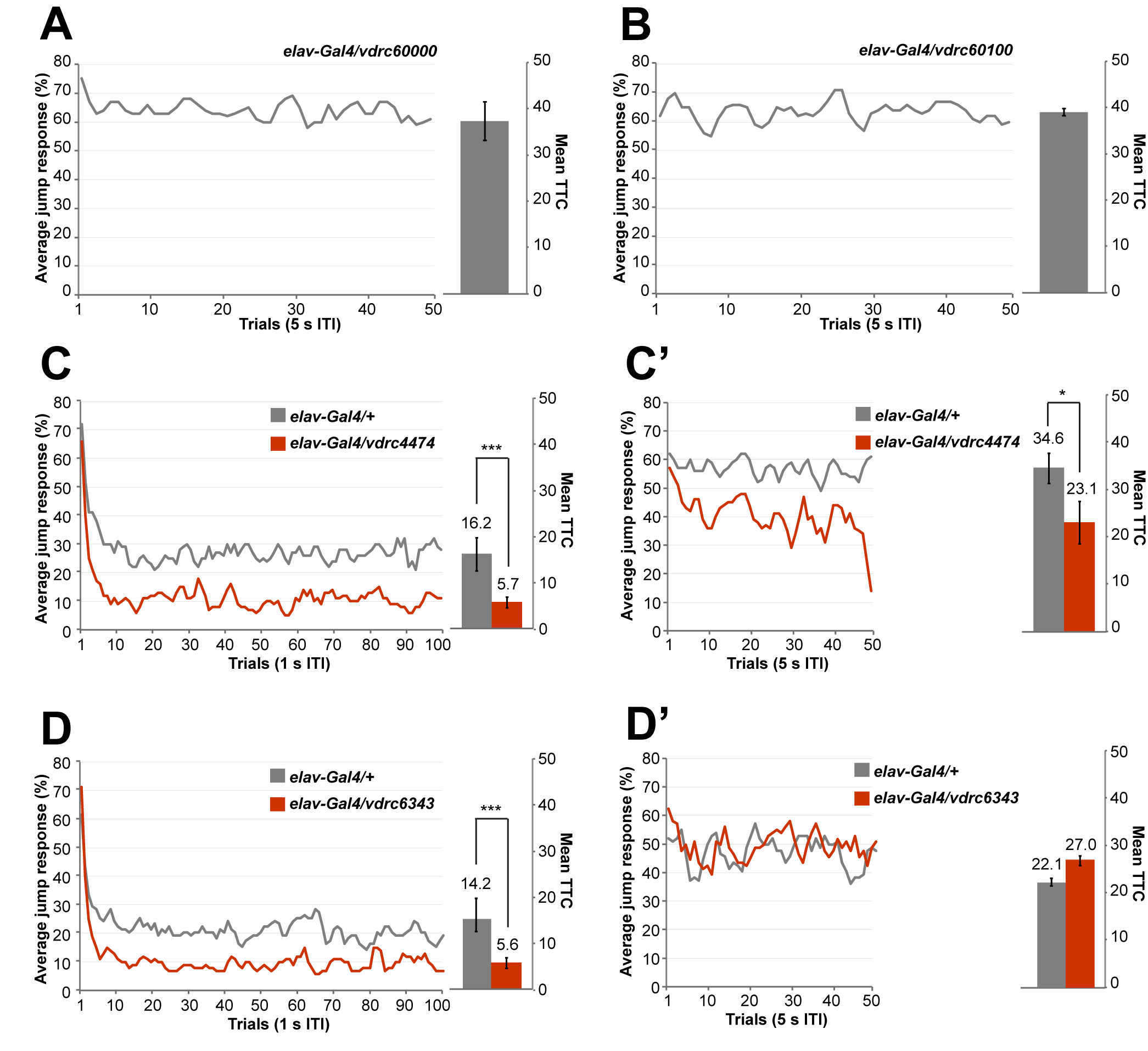

**Figure S4. Jump responses in the fatigue assay**

In fatigue assay, jump responses were induced by repeated light-off pulses for 50 trials with 5 s inter-trial interval (ITI). The average jump response in each trial was scored as % of jumping flies. The mean number of trials that flies needed to reach the no-jump criterion in the fatigue assay was scored as mean trials to criterion (Mean TTC), presented as Mean TTC ± SEM. (A, B) Jump response of elav-Gal4/vdrc60000 control flies as well as elav-Gal4/vdrc60100 flies (2xGMR-wIR/+; elav-Gal4, UAS-Dicer-2/+) remains high throughout the entire course of the experiment (60-70% jumping flies, mean TTCvdrc60000=37.3, mean TTC vdrc60100=40.4), illustrating that an ITI of 5 s prevents habituation from occurring. (C-D’) Representative examples of genotypes with premature habituation phenotype, one with and one without fatigue. (C) Premature habituation induced by repeated light-off pulses with 1 s ITI of elav-Gal4/vdrc4474 RNAi knockdown (N=51, in red) compared to its respective genetic background control (N=77, in gray). (C’) Decline of jump response of elav-Gal4/vdrc4474 RNAi knockdown (N=43, in red) compared to its genetic background control (N=58, in gray) in the fatigue assay (5 s ITI) indicating that premature habituation is confounded by fatigue. (D) Premature habituation (1 s ITI) of elav-Gal4/vdrc6343 RNAi knockdown (N=64, in red) compared to its respective genetic background control (N=75, in gray). (D’) No decline of jump response of elav-Gal4/vdrc6343 RNAi knockdown (N=26, in red) compared to its genetic background control (N=25, in gray) in the fatigue assay (5 s ITI) indicating that fatigue is unlikely to account for the observed premature habituation phenotype. *** p<0.001, * p<0.05, based on lm analysis.

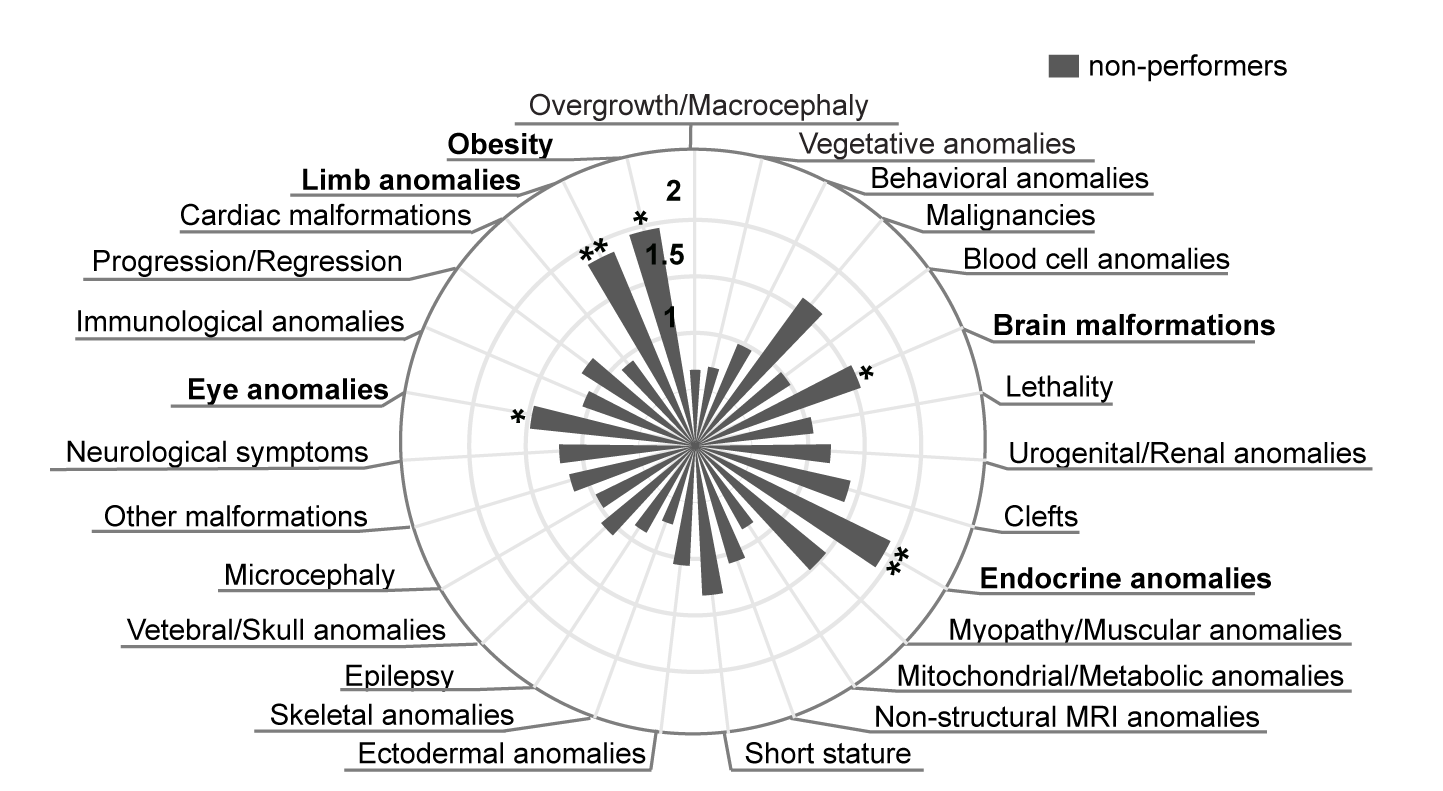

**Figure S5. Enrichment of genes associated with non-performers in ID subclasses and ID-associated clinical core features**

Genes belonging to the "non-performers" category show specificity for disorders accompanied by endocrine anomalies (E=1.9; 0.001), limb anomalies (E=1.86; p=0.008), eye anomalies (E=1.47; p=0.022), brain malformations (E=1.55; p=0.043) and obesity (E=1.95; p=0.015). ** p<0.01, * p<0.05 based on Fisher’s exact test. A complete list of enrichment scores and p-values is in Table S4.

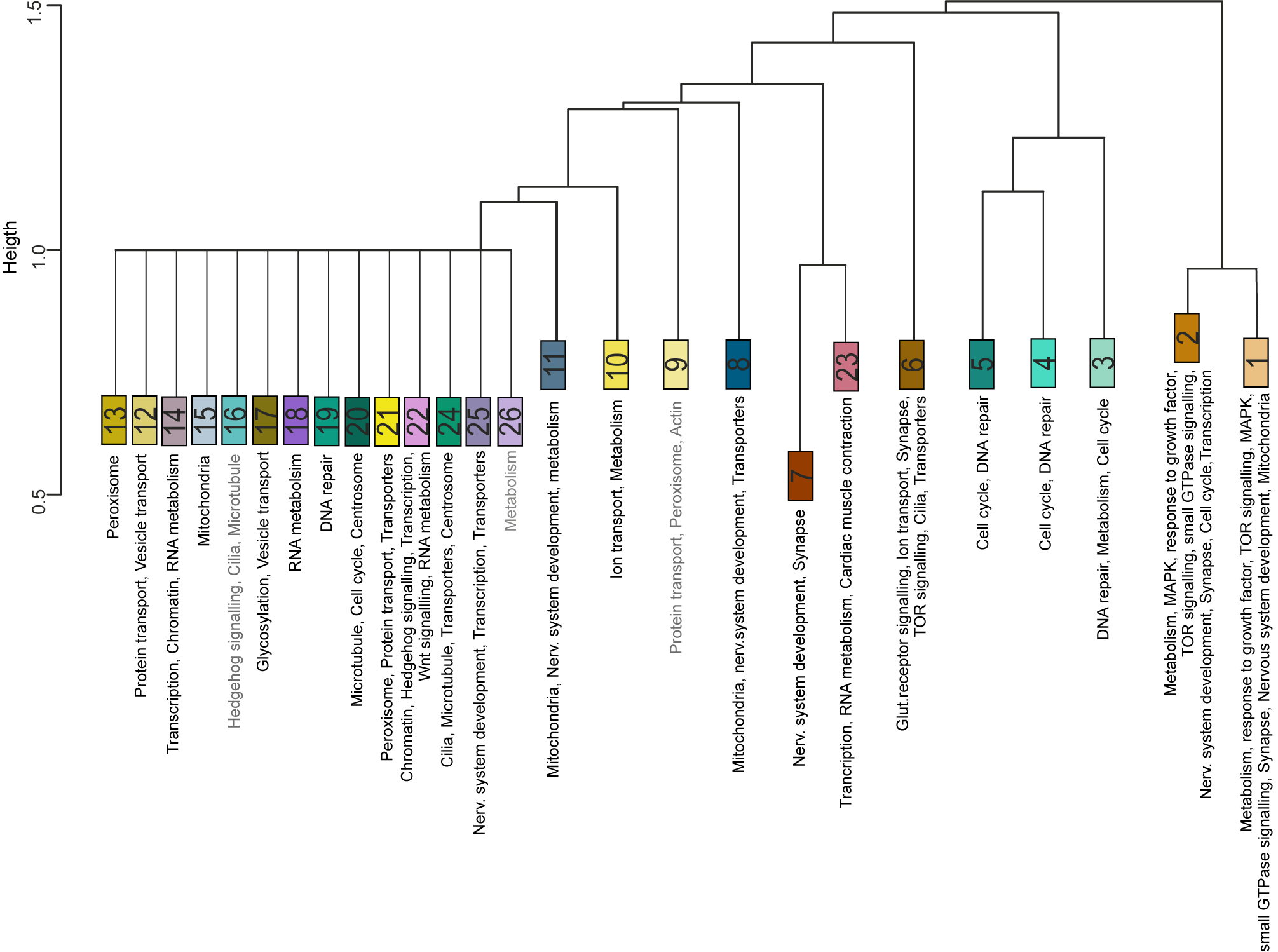

**Figure S6. Clustering of protein-protein interaction communities**

Interaction communities hierarchically clustered based on the numbers of nodes shared by pairs of communities. The enriched GO-based annotations are depicted. Communities are colored according to their GO proximity

**Tables:**

**Table S1**

Catalog of investigated human ID genes extracted from the SysID database (sysid.cmbi.umcn.nl), category primary ID genes (provided as a separate Excel sheet)

The table contains the following columns:

**Human ID gene symbol**

**Entrez ID**

**SysID** (from SysID database, sysid.cmbi.umcn.nl)

**Ensembl ID**

**Proposed/predicted disease mechanism** (see also **Proposed disease mechanisms**)

**Syndromic** (YES/NO; from SysID database, sysid.cmbi.umcn.nl)

**Non-syndromic** (YES/NO; from SysID database, sysid.cmbi.umcn.nl)

**ASD SSC** (YES/NO)

**ASD SFARI** (YES/NO)

**Table S2**

Catalog of investigated human ID genes, corresponding *Drosophila* orthologs, RNAi lines, and habituation results (provided as a separate Excel sheet).

The table contains the following columns:

**Human gene info**: Human ID gene symbol

**Fly gene info**: CG number, Flybase ID, fly gene symbol

**RNAi/vdrc: Ordernumber**

**Habituation screen results**: Lethality (YES/NO), Validated by independent repeat (YES/NO, if more than one experiment was done; see also **Phenotype reproducibility in the light-off jump habituation screen**), N=number of jumpers, % of jumpers (from total N), Mean TTC fold-change (calculated as Mean TTC_mutant_/Mean TTC_control_)

**lm analysis results**: Estimate, St. error, p-value (see also **Statistical analysis**)

**FDR correction**: p-value adjusted (see also also **Statistical analysis**)

**Fatigue phenotype**: Fatigue/No fatigue/Not tested (only genotypes with premature habituation were tested for fatigue)

**Phenotype category/RNAi line**: not affected/habituation deficit/non-performers/premature habituation

**Phenotype category/human ID gene**: not affected (0/1), habituation deficient (0/1), non-performers (0/1), premature habituation (0/1)

**Table S3**

Distribution of human ID genes in four phenotype categories identified in *Drosophila* light-off jump habituation screen (provided as a separate Excel sheet)

**Table S4**

Enrichments (provided as a separate Excel sheet)

**Table S5**

Catalog of investigated SSC and SFARI genes (provided as separate Excel sheet)

**Table S6**

MANOVA comparisons of quantitative measures of behavior and cognition in SSC

| **Trait** | **Habituation deficient** | | **Not affected** | | **F-statistics** | **p-value** |
| --- | --- | --- | --- | --- | --- | --- |
|  | **Mean** | **SD** | **Mean** | **SD** |  |  |
| IQ | 73.36 | 25.94 | 68.58 | 26.00 | 0.26 | 0.61 |
| SRS | 80.55 | 13.20 | 82.67 | 7.95 | 0.26 | 0.62 |
| CBCL-Int | 60.27 | 9.95 | 58.50 | 6.92 | 0.30 | 0.59 |
| CBCL-Ext | 60.32 | 11.77 | 57.00 | 5.86 | 0.83 | 0.37 |
| ABC | 53.64 | 24.92 | 35.92 | 21.07 | 4.35 | 0.04 |
| inappropriate speech | 5.45 | 3.16 | 1.42 | 1.68 | 16.85 | 0.00026 |
| hyperactivity | 19.27 | 11.89 | 12.33 | 8.75 | 3.14 | 0.86 |
| irritability | 13.68 | 8.91 | 8.75 | 6.51 | 2.83 | 0.10 |
| lethargy | 11.77 | 8.46 | 9.83 | 9.36 | 0.38 | 0.54 |
| stereotypy | 3.45 | 3.45 | 3.58 | 3.70 | 0.01 | 0.91 |

**Table S7**

Protein-protein interaction communities (provided as a separate Excel sheet)

**Table S8**

Previously identified ID/ASD genes with established habituation deficit

| **Habituation to: startle stimulus, novel object, novel environment, novel odor, stress** | | | | | | | |
| --- | --- | --- | --- | --- | --- | --- | --- |
| **Human gene** | **Organism** | **Ortholog name** | **Source** | **Downloaded** | **Previously studied as disease model?** | **Tested in present study?** | **Habituation deficit in**  **present study?** |
| NF1  PC  SLC12A5  CCDC88A  SHANK3  THRB  ATP1A3  L2HGDH  SYN1  GATAD2B  EHMT1  PDE4D  FMR1  GRIN1  FMR1  PARK2  DISC1  CAMTA1  CNTNAP2  CASK | zebrafish  zebrafish  mouse  mouse  mouse  mouse  mouse  mouse  *Drosophila*  *Drosophila*  *Drosophila*  *Drosophila*  *Drosophila*  rat  mouse  mouse  mouse  *Drosophila*  *Drosophila*  *Drosophila* | nf1a  pcxa  Slc12a5  Ccdc88a  Shank3  Thrb  Atp1a3  L2hgdh  Syn  simj  G9a  dnc  Fmr1  Grin1  Fmr1  Park2  Disc1  Camta  Nrx-IV  CASK | PhenomicDB  PhenomicDB  PhenomicDB  PhenomicDB  PhenomicDB  PhenomicDB  PhenomicDB  MGI  Flybase  Flybase  Flybase  Flybase  Flybase  (1)  (2)  (3)  (4)  (5)  (5)  (6) | 08122016  08122016  08122016  08122016  08122016  08122016  08122016  08122016  08122016  08122016  08122016  08122016  08122016  NA  NA  NA  NA  NA  NA  NA | Yes  No  No  No  Yes  Yes  Yes  Yes  No  Yes  Yes  No  Yes  Yes  Yes  Yes  Yes  No  No  No | Yes  Yes  No  No  Yes  No  No  Yes  Yes  Yes  Yes  Yes  Yes  Yes  Yes  No  No  No  Yes  Yes | Yes  Yes  No  No  Yes  No  No  No  Yes  Yes  Yes  Yes  Yes  No  Yes  No  No  No  No  Yes |

**Supplemental Methods:**

**SysID inclusion criteria**

All investigated genes are from the ‘primary ID gene’ category of the SysID database (sysid.cmbi.umcn.nl) (7). These genes contain mutations which were published as causative for intellectual disability with robust clinical evidence (e.g. a minimum number of three independent patients with *de novo* mutations and information on clinical features and cognitive abilities) and causative character of the mutations (based on number and nature of variants, segregation).

**In/exclusion criteria of experimentally investigated genes**

The vast majority, 252 genes, are extracted from the first data freeze of the SysID database (status mid 2010, ID data freeze 388). Of the 388 genes contained in this data freeze, these represent i) all genes with a (one-to-one or many-to-one) ortholog in fly (see below) ii) for which RNAi lines were available and iii) which were not completely lethal upon panneuronal knockdown, precluding any behavioral assessment. Habituation data from recently published (8–12) and unpublished projects focused on novel ID genes identified in our clinic or by external collaborators were included in the present study if carried out under the utilized standard conditions. Genes from these projects belong to later data freezes of the SysID database. The catalog of investigated genes, the data freeze to which they belong, and distribution in syndromic and non-syndromic ID disorders is provided in **Table S1** (SysID data freeze group).

***Drosophila* ID gene orthologs**

Orthologs of the 286 investigated human ID genes were mapped in ENSEMBL (Ensemblv72_June2013) (13), and treefam (14) annotations, and curated manually as described in Kochinke et al. (7). One-to-one and one-to-many (fly-to-human) criteria identified 316 orthologs for 422 human ID genes (ID data freeze 388 plus 34 from later data freezes).

**Proposed disease mechanisms**

Loss-of-function (LOF) was proposed as a disease mechanism in case of recessive inheritance in general, and for X-linked and dominant variants when variant nature pointed to it (e.g. truncating) or when experimental support for (partial) loss-of-function as the underlying mechanisms was available (Pubmed, summarized on OMIM, the Online Mendelian Inheritance in Men database). This was the case for 262 of the investigated genes, corresponding to 92% of investigated ID/ASD disorders. In a few cases (5/262) supporting evidence for both LOF and gain-of-function (GOF) mechanisms has been reported. For mutations in 16 of the investigated genes, GOF mechanisms have been reported (<6%). The majority of these (9 genes) represent the core Ras-MAPK pathway. GOF of this pathway was appropriately modeled in our follow-up work on the screen. Lastly, for mutations in 8 genes no data was available that would allow conclusions about LOF versus GOF (<3%). Information about the proposed disease mechanism per gene is provided in **Table S1**. We concluded that in a large-scale approach, RNAi is a reasonable approximation to appropriately model the vast majority of studied ID/ASD conditions. ID genes with reported GOF mechanisms, because not recapitulated by our RNAi screen, were excluded from enrichment analysis of ID-associated clinical features, from enrichment analyses of genes implicated in ASD and from comparison of quantitative measure of behavior and cognition in ASD SSC.

***Drosophila* stocks, crosses and maintenance**

*Drosophila* stocks and crosses were reared on a standard *Drosophila* diet and raised at 25°C/70% humidity. Tissue-specific knockdown was achieved with the UAS-Gal4 system (15). In-house generated or assembled Gal4 lines were: *w1118; 2xGMR-wIR; elav-Gal4, UAS-Dicer-2* (panneuronal), *w1118; Gad1-Gal4; 2xGMR-wIR* (GABAergic), *w1118; Cha-Gal4, UAS-GFP; 2xGMR-wIR* (cholinergic). The *2xGMR-wIR* element suppress pigmentation in the eye, as required for an efficient light-off jump response (8, 9). UAS-Dicer-2 was used in the panneuronal Gal4 driver line to increase RNAi efficiency (16). UAS-RNAi lines and their genetic background controls (#60000 for RNAi lines from GD library and #61000 for RNAi lines from KK library) were obtained from the Vienna *Drosophila* Resource Centre (VDRC, www.vdrc.at). The *UAS-Raf^GOF^* line was obtained from Bloomington *Drosophila* Stock Center (#2033). *Ras1^R68Q^* and *UAS-Ras1^R68Q^* lines were kindly provided by H. Steller (17). These were crossed into the VDRC genetic background #60000 for seven generations and compared to this isogenic background. Knockdown or overexpression was induced by crossing Gal4 driver and transgenic UAS lines. Progeny of crosses between Gal4 driver and the corresponding genetic background of GD or KK libraries served as daily controls in all experiments. Male progeny (or female virgin progeny for UAS-RNAi insertions on the X chromosome) of appropriate genotypes was tested.

**Phenotype reproducibility in the light-off jump habituation screen**

To evaluate phenotype reproducibility in our screen and estimate false positive and false negative rates, all genotypes with significant effect on log TTC (p>0.05 at n=32) plus 100 randomly selected genotypes without significant effect on log TTC in light-off jump habituation (p≥0.05) were subjected to independent repeats. An additional 32 flies per genotype, plus an additional 32 control flies were tested. Based on 98% of confirmed significant effects, the false positive rate was estimated to be 2%. Genotypes without significant effect did not show any difference from the first testing, estimating the false negative ratio due to experimental variation to be near 0%. False negatives due to incomplete efficiency of the RNAi construct cannot be excluded. Results from the independent repeats were combined with the results of the systematic screen and are shown in **Table S2** (see also **Statistical analysis**).

**Validation of the wing engagement in the light-off jump response in tethered flies**

For high-speed, high-resolution videos of fly responses to a light-off stimulus, experiments were performed using a modified tethered escape assay (18). In brief, flies were cold anesthetized in an ice bath before being transferred to a cold plate for mounting onto tethers. Flies were allowed to recover for 30 minutes prior to being enclosed within the assay, where they were acclimated for 10 minutes before the start of each experiment. A flightstop arm was engaged continuously during both acclimation and trial presentation to stop flight after it was initiated. The projection screen used for the induction of looming stimulus, as described in Goodman et al. (18), was substituted with an LED array recapitulating the technical parameters of the LED illumination in the vials of the light-off jump habituation system. The LED array consisted of a 10x40 mm array of 7 evenly spaced LEDs covered by a white diffusor (30% transmittance). The luminance measured at the location of the fly, 6mm from the diffusor, was 5.3 Lux, generated by a 0.0149 mA current through the array. Each fly was subjected to a light off stimulus (15ms). The videos were recorded at a 256x256 resolution (approximately 67 pixels/mm scale) at 3000 frames per second. **Video S1** (right) shows the engagement of the wing vibration in this type of escape response.

**Validation of the automated jump scoring and of jump specificity**

The automated scoring of jump responses by audio recording of wing vibrations was compared to manual jump scoring in simultaneous audio and video recordings, first for a set of controls and subsequently for three habituation deficient and for three not affected habituation genotypes selected from the screen. In a regular habituation experiment, i.e. 32 flies exposed to a series of 100 light-off pulses with 1 s interval, responses in 93.5% of trials were identical between the video and audio, in 4% of trials the jump responses were only detected in the video and in 2.5% of trials the jump responses were only detected by audio recordings. Thus, the estimated rate of missed jumps is 4% and the estimated rate of false positive jumps is 2.5%. The effect of a gene knockdown on habituation was assessed by manual scoring of TTC values, analyzed vial by vial. Both video and audio scoring, when performed simultaneously, was able to faithfully recapitulate the phenotypes (**Figure S1**). In the screen, flies of all genotypes were monitored before and during the experiment. None of them was found to show excessive jumping/jumping without stimulation, excessive locomotion or affected climbing behavior after tapping them down to the bottom of the vials in which they were housed prior to being filled into the light-off jump habituation chambers. They jump with precision in response to light-off stimuli, within the first 100 ms following the stimulus (**Videos S2,** **S3** and **Figures S2, S3A**). Along with habituation, reactivity to slightly dimmed light (down to 75% of default intensity) was measured, which in controls and all genotypes did not effectively elicit jump response, excluding hypersensitivity (**Figure S3B**).

**Fatigue assay**

Fatigue was tested for each genotype that resulted in premature habituation (for data see **Table S2**, column O: Fatigue/No fatigue/Not tested). The fatigue assay was performed and analyzed in the same way as the light-off jump habituation assay, with the following adjustments. The interval between light-off pulses was increased to 5 seconds, an intertrial interval that is sufficiently long to prevent habituation/formation of non-associative learning in this assay. The light-off pulse was repeated 50 times. Control flies did not reduce their jump response after 50 trials of light-off stimulus in the fatigue assay (**Figure S4**). Fatigue was concluded when log-transformed TTC values of the mutant were significantly smaller than log-transformed TTC values of the control (non-corrected p-value < 0.05; based on the utilized linear model, see **Statistical analysis**).

**Quality criteria for RNAi lines**

To minimize the number of potential off-targets, 98% of all constructs used in vdrc UAS-RNAi lines had an s19 score > 0.98, a previously established quality criteria (7, 19). No effect on viability, jump response, or habituation was seen when crossing the panneuronal Gal4 driver to the ‘40DUAS’ line (20), containing UAS repeats but no functional short hairpin RNA (a potential source for dominant phenotypes due to an integration locus of the VDRC KK library (20, 21).

**Annotation of ID plus ASD-associated genes**

ID genes represented in our screen that carried LGD mutations (22) and mutations predicted as (probably and possibly) damaging (Table S5, Sanders et al., 2015) (23) in ASD individuals of SSC families, a well-phenotyped cohort of families with one affected individual (24), were extracted and identified as ID plus ASD-associated genes (ASD SSC, 33 genes). ID genes represented in our screen and listed in the high confident categories (S = syndromic, 1 = high confidence, 2 = strong candidate, 3 = suggestive evidence) of the SFARI database ([www.sfari.org](http://www.sfari.org), update July 10, 2017), a gene reference resource for ASD research (25, 26), were extracted and identified as ID plus ASD-associated genes (ASD SFARI, 38 genes). Both were used for subsequent enrichment analyses. Genes are listed in **Table S5**. The distribution of the investigated ID genes in ASD SSC and ASD SFARI categories is indicated in **Table S1**.

**Identification of enriched Gene Ontology-based (GO) categories**

32 GO-based categories previously reported to be enriched in ID (7) were analyzed for enrichment among the genes investigated in the present study as previously described (7). Out of these, 25 GO categories were also enriched among the investigated 286 ID genes investigated in this study (**Figure 2**). Enrichment of these 25 GO-based categories among habituation phenotype-based gene groups was determined.

**Enrichment analysis**

The enrichment of human ID-accompanying clinical features, ASD-associated genes, GO-based categories and *Drosophila* datasets among habituation phenotype-based gene groups were calculated as follows: (a/b)/((c-a)/(d-b)), whereby a=genes in ID-accompanying clinical feature/ID plus ASD group/GO-based category and with investigated habituation phenotype, b=genes in ID-accompanying clinical feature/ ID plus ASD group/GO-based category, c=genes with investigated habituation phenotype, d=background/all genes in screen.

**Comparison of quantitative measure of behavior and cognition in ASD SSC**

Scores for following five broad quantitative measures were extracted for ASD SSC individuals with LGD mutations and predicted damaging mutations in genes associated with habituation deficit and genes with no habituation deficit in *Drosophila*: measurement of cognitive ability (full-scale IQ, IQ); ASD-associated behaviors (Social Responsiveness Scale, SRS (27)); symptoms associated with depression and anxiety (Child Behavior Checklist Internalizing Disorders, CBCL-Int (28)); symptoms associated with impulsivity, attention and conduct (Child Behavior Checklist Externalizing Disorders, CBCL-Ext (28)), and atypical behavior (Aberrant Behavior Checklist, ABC (29))). MANOVA comparisons between “habituation deficient” and “not affected” groups were performed for all five measures and for five subdomains of ABC: inappropriate speech, hyperactivity, irritability, lethargy and stereotypy. Complete results of MANOVA comparisons are listed in **Table S6**.

**Molecular interaction network**

For analysis of molecular interactions, we created a human-specific interaction network containing three categories of interactions: (1) physical, including predicted interactions from other species (source: GeneMANIA (30) – release Jul 10, 2014; HPRD, Human Protein Reference Database (31) – release 9, Apr 13, 2010), (2) shared protein domains (source: GeneMANIA – release Jul 10, 2014), and (3) pathway interactions (source: GeneMANIA – release Jul 10, 2014). This reference network was used for community clustering and enrichments.

**Community clustering**

Interaction communities were obtained from the R package ‘linkcomm’ (32) and hierarchically clustered based on the number of shared nodes (after scoring the pair-wise similarities using the Jaccard coefficient; tree cut-off at 0.9).

**PIE score**

The physical interaction enrichment (PIE) score and associated p-value were calculated against interactions from the reference network, using the PIE algorithm (33) to account for biases in the number of reported interactions as frequently associated with disease-associated genes. Random protein groups were formed by number-matched sub-samplings selected from all interactions in the reference interaction network.

**Data visualization**

Graphs were generated with Microsoft Excel. Polar bar graphs in Figure 2A, Figure 4 and Figure S4 were generated with JS Highcharts (www.highcharts.com). Cluster dendrogram in Figure S3 was generated with R (34). Network visualization in Figure 5A was carried out using Cytoscape (v 2.8.3 (35)).

**Statistical analysis**

Statistical analyses were performed using SPSS and R statistical software (34). No statistical analysis was used to predetermine sample size. The distribution of data was examined with histogram. Data with non-normal distribution were log transformed. Sample sizes, statistical tests and p-values are indicated in the text, figures, figure legends and supplemental tables. Main effects of the genotype on log-transformed TTC values in the light-off jump habituation assay and in the fatigue assay were tested using a linear model regression analysis (lm) in the R statistical software (34)) and corrected for the effects of testing day and system. Final p-values were calculated based on independent experimental repeats and corrected for false discovery rate (FDR) using Benjamini-Hochberg correction (36). Corrected p-values < 0.05 were considered significant (p_adj_). In the enrichment analyses, uncorrected p-values were determined using two-sided Fisher’s exact test in R (34). MANOVA (SPSS) was used for comparison of quantitative measure of behavior and cognition in ASD SSC. A stricter p-value was applied to determine significance in case of multiple comparisons (5 comparisons: p< 0.01).
